# Iron Regulatory Protein 1 is Required for the Propagation of Inflammation in Inflammatory Bowel Disease

**DOI:** 10.1101/2023.01.27.525690

**Authors:** L. Fahoum, S. Belisowski, N. Ghatpande, N. Guttmann-Raviv, W. Zhang, K. Li, W-H. Tong, A. Nyska, M. Waterman, R. Weisshof, A. Zuckerman, E.G. Meyron-Holtz

**Affiliations:** Laboratory of Molecular Nutrition, Department of Biotechnology and Food Engineering, Technion– Israel Institute of Technology, Haifa, Israel; Jiangsu Key Laboratory of Molecular Medicine, Medical School of Nanjing University, Nanjing 210093, China; Molecular Medicine Program, Eunice Kennedy Shriver National Institute of Child Health and Human Development, Bethesda, MD 20892, USA; Tel Aviv University and Consultant in Toxicologic Pathology, Tel Aviv, Israel; Rambam / Technion– Israel Institute of Technology, Haifa, Israel; Aviv Projects, Ness Ziona, Israel

**Author notes:** These authors both contributed most significantly to this work (L. Fahoum, S. Belisowski).

## Abstract

**Objective:** Inflammatory bowel diseases (IBD) are complex disorders. Iron accumulates in the inflamed tissue of IBD patients, yet neither a mechanism for the accumulation nor its implication on the course of inflammation are known. We hypothesized that the inflammation modifies iron homeostasis, affects tissue iron distribution and that this in turn perpetuates the inflammation.

**Design:** This study analyzed human biopsies, animal models and cellular systems to decipher the role of iron homeostasis in IBD.

**Results:** We found inflammation-mediated modifications of iron distribution, and iron-decoupled activation of the iron regulatory protein (IRP)1. To understand the role of IRP1 in the course of this inflammation-associated iron pattern, a novel cellular co-culture model was established, that replicated the iron-pattern observed in vivo, and supported involvement of nitric oxide in the activation of IRP1 and the typical iron pattern in inflammation. Importantly, deletion of IRP1 from an IBD mouse model completely abolished both, the misdistribution of iron and intestinal inflammation.

**Conclusion:** These findings suggest that IRP1 plays a central role in the coordination of the inflammatory response in the intestinal mucosa and that it is a viable candidate for therapeutic intervention in IBD.

**What is already known on this topic:** Inflammatory bowel diseases (IBD) are chronic conditions that cause inflammation in the digestive tract. Iron accumulation is a common feature of IBD, but the mechanism of accumulation and its implications for the course of inflammation are not fully understood.

**What this study adds:** This study reveals an inflammatory intestinal iron distribution-pattern, that has not been described previously involving iron accumulation in immune cells and iron deficiency in epithelial cells. We show that an inflammation mediated activation of the Iron Regulatory Protein (IRP)1 is responsible for this inflammatory iron pattern and that this drives the propagation of the inflammation in IBD. Moreover, targeted deletion of IRP1 completely abolished the intestinal inflammation.

**How this study might affect research, practice or policy:** These findings suggest that IRP1 plays a central role in the coordination of the inflammatory response in the intestinal mucosa in IBD. This might lead to the development of novel treatment approaches for IBD focused on modulating IRP1 activity.

## Introduction

Inflammatory bowel diseases (IBDs) are long-term multifactorial inflammations of the gastrointestinal tract, with Crohn’s disease (CD) and colitis being the two most common forms. Since the beginning of industrialization, global incidence has been rising and due to a lack of cure, global prevalence increases as well.^1^ Although the cellular and molecular events initiating and perpetuating the inflammation are not fully understood, it is clear that intestinal homeostasis is broadly disrupted, causing pathological inflammation that leads to tissue injury. Increased iron concentrations have been measured in the inflamed tissues of Crohn’s disease and colitis patients, when compared to tissue of healthy individuals and to patient samples collected from noninflamed regions,^2^ but neither the impact of, nor the mechanism driving this phenomenon are known.

Iron is an essential nutrient, yet unbalanced iron levels lead to inflammation and tissue damage.^3–5^ Iron homeostasis is tightly regulated at the systemic and cellular level, primarily by the peptide hormone hepcidin^6^ and by the two iron regulatory proteins (IRP1/2) respectively.^7^ Hepcidin acts by binding to the iron exporter ferroportin, where it sterically inhibits iron export and causes ferroportin degradation.^8,9^ IRPs regulate numerous transcripts related to cellular iron import, storage and export, by binding with high affinity to RNA motifs known as iron-responsive elements (IREs). For example, binding of IRPs to the IRE in the 5’ untranslated region (UTR) of the subunits (FTH1 and FTL) of the iron storage protein ferritin,^10^ inhibits their translation. Reducing the cytosolic ferritin levels will lead to less storage capacity and elevate iron availability. In contrast, IRP binding to the iron importer transferrin receptor (TFR1) 3’ UTR IREs, stabilizes these transcripts, leading to increased iron import.

IRP1 is a bifunctional protein that primarily functioning as a cytosolic, iron-sulfur cluster-containing, aconitase (cAco). In iron deficient conditions, it undergoes a conformational change and turns into an RNA (IRE)-binding apoprotein. Similar to IRP1, IRP2 binds to IREs in iron deficient conditions, yet it is degraded in the presence of high iron. Both IRPs are also regulated by oxygen and reactive oxygen- and nitrogen-species (ROS/RNS). However, in contrast to their concerted iron regulation, their response to oxygen is opposing.^11,12^ IRP2 is stable at physiologic oxygen and acts as an RNA-binding protein under these conditions. In contrast, the Fe-S cluster of cAco is relatively stable under physiologic oxygen conditions, therefore IRP1 maintains its aconitase activity, even when iron levels are low.^13–15^

Similar to the opposing response of the two IRPs to oxygen, they also respond differently to nitric oxide (NO). NO plays important roles in pathogen killing, as well as macrophage metabolic and epigenetic remodeling, and is elevated in inflammatory processes including IBD.^16^ The biological consequences of NO-mediated IRP1 activation versus IRP2 degradation on cellular iron, TfR1 and ferritin, were extensively studied in macrophages,^17^ yet their systemic role in specific inflammatory diseases remains unclear. We hypothesized that NO-mediated IRP1 activation drives the observed iron accumulation in the inflamed tissues of IBD patients, and thus perpetuates the inflammation.

The present work studied iron homeostasis in biopsies from CD patients, animal models for CD and colitis and a cellular co-culture model that mimics the interaction between intestinal epithelium and immune cells. IRP1 was found to be activated by inflammation. This inflammation mediated regulation of IRP1 activity impaired intestinal iron homeostasis and arbitrated the propagation of inflammation in IBD. Most importantly, targeted deletion of IRP1 minimized the inflammation, flagging it as a novel target for the treatment of IBD.

## Results

### Intestinal inflammation triggers an iron independent IRP1 activation and a cell specific pattern of iron distribution

Tumor necrosis factor alpha-overexpressing mice (*Tnf* ^*ΔARE/+*^), which are a genetic mouse model for CD,^18^ and the chemical DSS model for colitis^19^ were used to evaluate the effect of inflammation on iron homeostasis and IRP activity. *Tnf* ^*ΔARE/+*^ mice develop spontaneous inflammation in the terminal ileum early and rheumatoid arthritis later in life.^18^ Intestinal sections of 12-14-week-old *Tnf* ^*ΔARE/+*^ mice were stained for ferritin, a sensitive indicator of cellular iron status.^20^ While ferritin accumulated in the immune cells, its levels were significantly reduced in intestinal epithelial cells (IECs) of the inflamed terminal ileum of *Tnf* ^*ΔARE/+*^ mice compared to wild-type (WT) controls (Fig. 1A, Fig.S1A). Similar trends were observed in sections from mice with DSS-induced colitis (Fig. 1A, Fig. S1A). To test if other iron-related proteins respond similarly, the terminal ileum of *Tnf* ^*ΔARE/+*^ mice was fractionated into an IEC- and an immune cell-enriched (lamina propria-LP) fraction. Various proteins, known to respond to cellular iron levels, were analyzed in each fraction separately. Consistent with the ferritin response to inflammatory conditions, IEC TFR1 levels were increased and LP TFR1 levels reduced, when compared to the WT fractions (Fig.1B). In addition, IRP2 levels were increased in the inflamed IECs and decreased in the inflamed LP (Fig. 1C). In contrast, IRP1-IRE-binding activity was significantly elevated in both fractions of the inflamed ileum compared to uninflamed ileum from WT mice, suggesting that the regulation of IRP1 activity is independent of the intracellular iron status (Fig. 1D). Taken together, intestinal inflammation triggered a unique pattern of iron distribution, involving iron accumulation in immune cells and iron deficiency in IECs.

**Figure 1:**
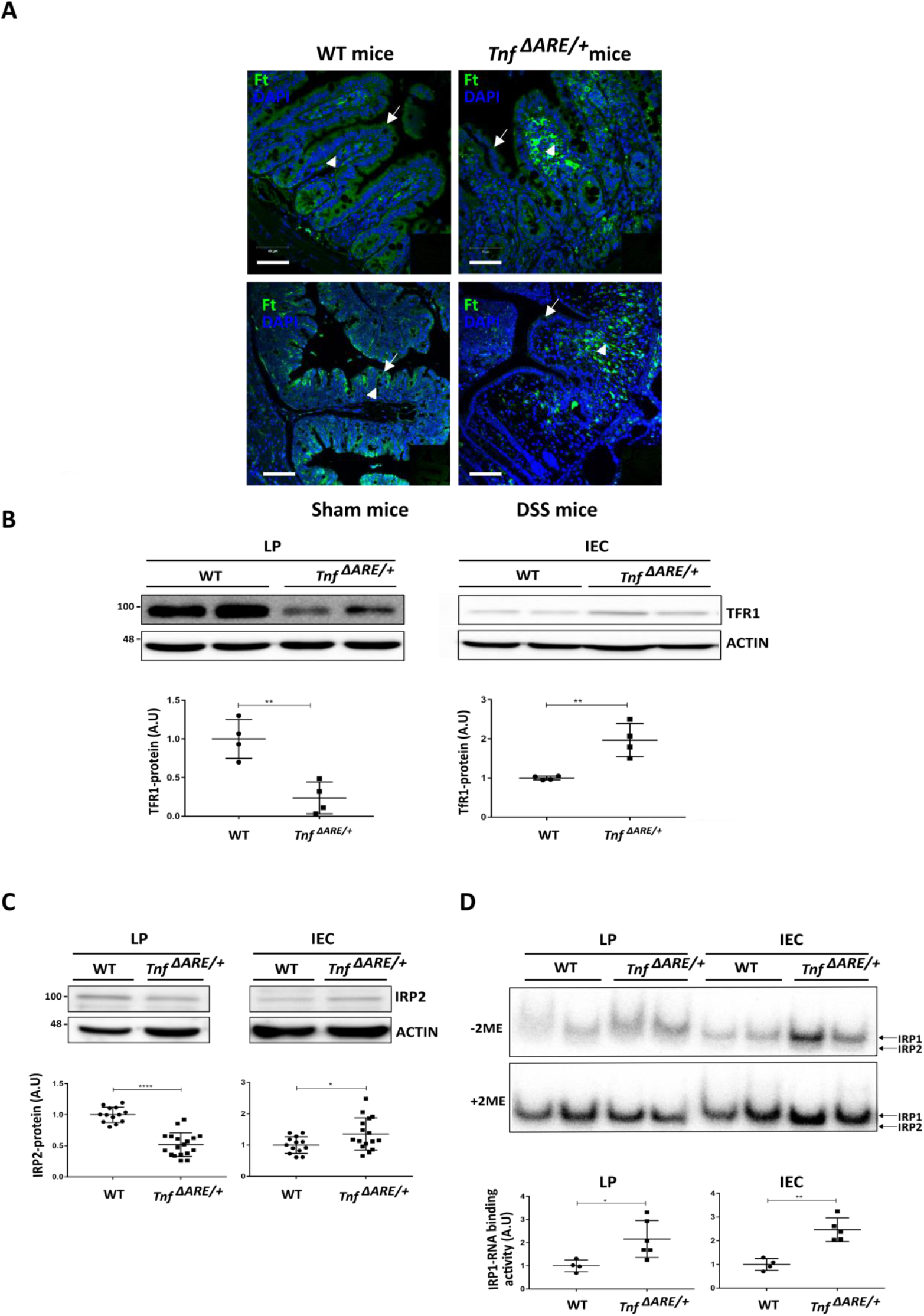
Inflammation drives a cell specific pattern of iron distribution. (**A**) Sections from terminal ileum (TI) of 12-14-week-old wild type (WT) and *Tnf* ^*ΔARE/+*^ mice (n=3 each), as well as C57BL/6 mice treated for 7 days with drinking water supplemented with 4% dextran sulfate sodium (DSS), were immuno-stained for ferritin (green). Representative images show decreased ferritin levels in the inflamed intestinal epithelial cells (IEC) (arrows). Lamina propria (LP) cells show increased ferritin levels (arrow heads) in both inflammation models. Scale bars 50 μm. (**B-D)** TI was harvested from 12-14-week-old WT and *Tnf* ^*ΔARE/+*^ mice. IEC- and LP-enriched fractions were isolated and lysates further analyzed. **B)** Representative immuno-blot for transferrin receptor-1 (TFR1) (mean ± SD (n=4)). **C)** Representative immuno-blot for IRP2 (mean ± SD (n>16)). **D**) Representative radiograph of electromobility shift assay (EMSA) and quantification of IRP1 RNA-binding activity. Addition of 2-mercaptoethanol (2-ME) to the lysate removes all Fe-S clusters from cAco/IRP1 and thus shows total IRP1 RNA-binding capacity. Values were normalized to +2-ME (mean ± SD (n>4)). Quantifications (B-D) were done, using Image J.

### Physical contact is required between macrophages and epithelial cells to mimic the activation of IRP1-IRE binding activity in vitro

We searched for a suitable cell-model to elucidate the forces driving the iron-independent activation of IRP1 and the intestinal inflammation-specific iron distribution pattern observed in mice. Several co-culture models of epithelial cells (EC) with immune cells separated by filter inserts, have been successfully employed to reproduce various aspects of inflammation,^21^ yet none of these models could reproduce inflammation-specific iron patterns. We hypothesized that NO produced by macrophages diffuses and activates IRP1 in neighboring ECs, and that this activation mediates the inflammatory iron pattern. Thus, we tested co-cultures with direct cell-to-cell contact between ECs and macrophages. To enable differentiation between EC- and macrophage-derived proteins, a direct interspecies co-culture model was developed with human epithelial enterocyte-like Caco2 cells, and either murine primary bone marrow-derived macrophages (BMDM) or the murine macrophage-like cell line Raw 264.7.^22^ Lipopolysaccharide (LPS)-induced inflammation was validated by the activation of critical markers of signaling pathways involved in inflammation, including MAP kinase pathways and NFkB-p65 (Fig. S2A). In addition, downstream proinflammatory cytokines, including *IL8* and *ICAM1* in ECs (Fig. S2B) and *Tnfα, Icam1, Il1b* and *Il6* in macrophages (Fig. S2C), were all activated in both co-cultures and the time-course of the LPS-triggered inflammatory response was nearly identical to that reported for *in vivo* and *in vitro* models of LPS-induced inflammation.^23^

The LPS-triggered response was identified in both macrophage mono-cultures and macrophage-EC co-cultures, but not in EC mono-cultures (Fig. S3A), suggesting the need for macrophages to elicit the epithelial inflammatory response. Indeed, application of the NO donor S-nitro-N-acetyl-D, L-penicillamine (SNAP) elicited an inflammatory response in EC monocultures (Fig. S3B). Further, in SNAP-treated EC monocultures, intracellular iron was reduced and extracellular iron increased (Fig. S3C), supporting the notion that NO generated in macrophages (Fig S3A) contributes to the inflammatory iron pattern in ECs.

In parallel, the RNA binding activity of macrophage IRP1 was significantly increased after LPS stimulation (Fig. 2A lane 3 and 4), similar to the observation in the *Tnf* ^*ΔARE/+*^ mice (Fig. 1D). Yet, epithelial IRP1 could not be evaluated in a gel shift assay, as it migrated together with macrophage IRP2. Therefore, IRP1 aconitase activity was evaluated using the in-gel aconitase-assay, as decreased cAco-activity correlates with increased IRP1 RNA-binding-activity. Gel conditions were optimized to enable differentiation between human vs. mouse cAco-activity (Fig. 2B lane 1 and 4). A decrease in cAco-activity was observed in response to LPS treatment in both cell types, which is consistent with an increase in the RNA-binding-activity of IRP1. Moreover, LPS treatment led to a significant decrease in mitochondrial aconitase (mAco)-activity in both cell types, suggesting a metabolic shift to glycolysis, as previously reported.^24,25^ Taken together, the direct interspecies co-culture model developed here, demonstrated activation of a broad range of inflammatory signaling pathways upon LPS-induced inflammation. Most importantly, the cell-model replicated the increased IRP1 mRNA-binding activity as seen in vivo (Fig.1D).

**Figure 2:**
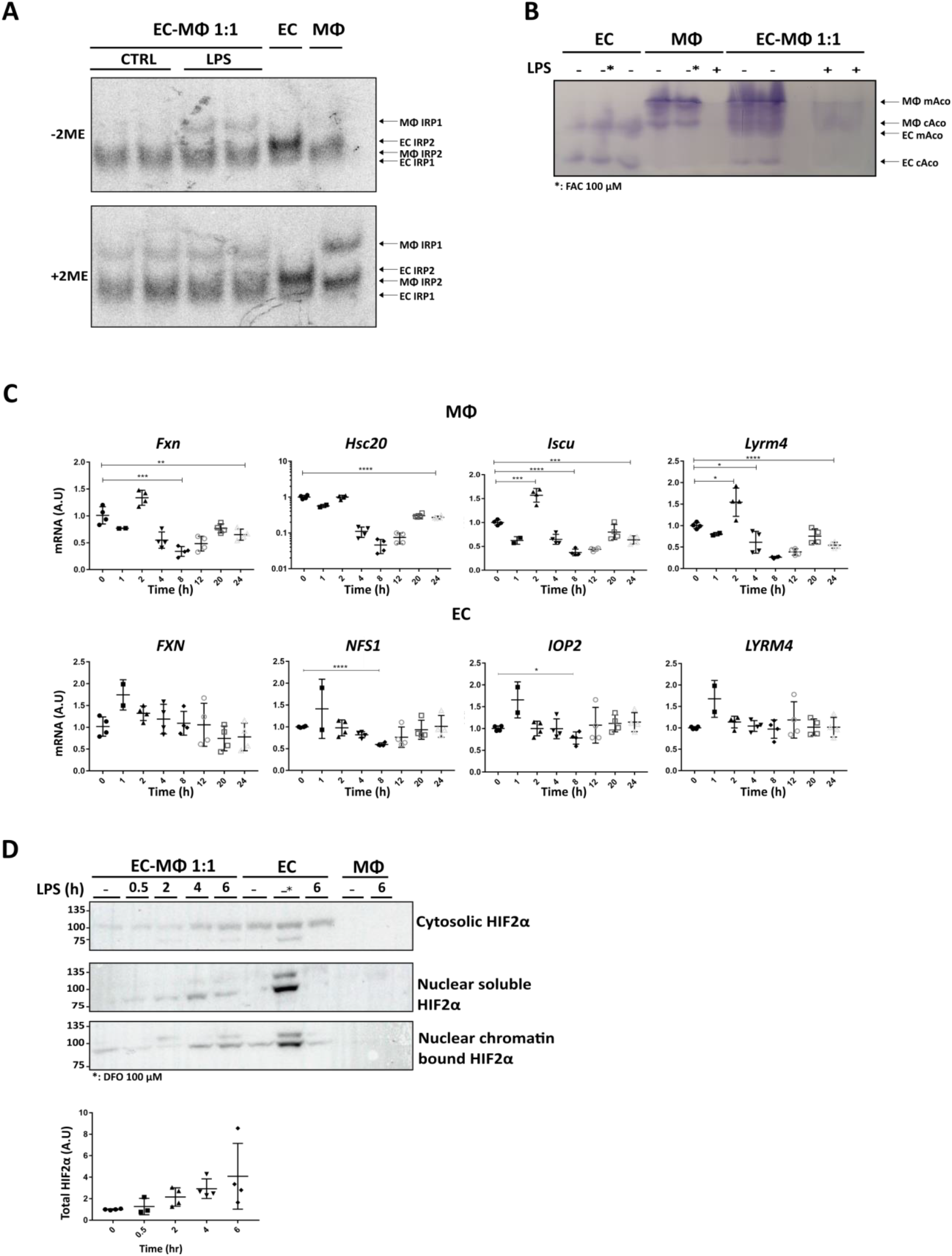
An interspecies co-culture model mimics the inflammation-associated activation of IRP1. Co-cultures of epithelial cells (EC) and macrophages (MΦ) were treated with 200 ng/ml lipopolysaccharide (LPS) for 24 h. Cells were harvested and lysed. (**A)** Lysates were subjected to electromobility shift assay (EMSA) for assessment of IRP-RNA binding activity (n= 4). A representative radiograph is shown. (**B)** Lysates were subjected to in-gel aconitase assay (n= 4). A representative gel is shown. **(C)** Expression levels of RNA of key proteins associated with iron-sulfur cluster biogenesis were measured by RT-qPCR. (**D)** Representative immunoblots of lysates after subcellular fractionation. Protein levels were quantified with Image J. Values are presented as mean ± SD (n=4). * 100 μM desferrioxamine (DFO) for 24 h. In co-cultures, EC were always Caco2 cells. MΦs were as follows: Raw264.7 in **A, B** and **D**, and bone marrow derived macrophages (BMDM) in **C**.

We reasoned that increased IRP1 RNA-binding activity may be due to impaired Fe-S cluster assembly. This was based on the fact that the cysteine desulfurase of the core iron-sulfur cluster assembly complex Nfs1 and the iron-sulfur cluster assembly enzyme IscU mRNA and protein levels were reported to be low in BMDM in response to inflammation.^26^ Thus, several genes involved in Fe-S cluster assembly were evaluated. In macrophages, the transcription levels of all four tested iron-sulfur cluster biogenesis proteins were significantly decreased, in agreement with the decrease in aconitase activities in mAco and cAco. However, in epithelial cells, transcripts of iron-sulfur cluster biogenesis proteins did not change (Fig. 2C). In IBD, the epithelial transcription regulator HIF2α is increased,^27^ and given that HIF2α is a key regulator of iron transport through epithelial cells,^28^ cytosolic and nuclear levels of HIF2α, were tested. We found that HIF2α was significantly increased in ECs, beginning 4 h after LPS stimulation (Fig. 2D). In line with this, 4 h after LPS stimulation, levels of both the apical iron importer DMT1 and the basolateral iron exporter FPN gradually increased (Fig. S4), likely promoting iron flux from the intestinal lumen, through the epithelial cells and into the blood. This can explain the observed epithelial iron deficiency and consequential increase in IRP1 RNA-binding activity. In summary, increased NO production, decreased iron-sulfur cluster assembly proteins in macrophages and increased HIF2α in epithelial cells support the inflammation-induced activation of IRP1-IRE binding activity in both cell types.

### The epithelial-macrophage co-culture mimics the cell specific pattern of iron distribution

To test the effect of inflammation on iron homeostasis, changes in ferritin-iron content were analyzed upon LPS stimulation of co-cultures. Epithelial ferritin iron was reduced and macrophage ferritin iron was elevated 24 h after LPS stimulation (Fig. 3A lanes 10 and 11). In addition, a strong increase in ferritin protein, especially of the FTH1 subunit, was detected in the macrophages of the co-cultures (Fig. 3B), consistent with the observations in macrophage mono-cultures.^29^ We thought, that the increase of ferritin in macrophages, despite the activation of macrophage IRP1, could be due to an inflammation-mediated decrease in ferritin degradation. Pulse-chase analysis of co-cultures that were metabolically labeled with ^35^S just before stimulation with LPS showed that macrophage ferritin subunit degradation was slower in LPS-treated co-cultures (Fig. 3C lane 9 and 11) compared to untreated controls (Fig. 3C lane 8 and 10). A decrease in ferritin degradation was further suggested by a decrease in transcript levels of the ferritinophagy-mediating *Ncoa4* in inflamed BMDM (Fig. 3D). After 24 h of LPS stimulation, macrophage ferritin migrated slightly faster in native gels (Fig. 3A lane 11), compared to its non-treated counterpart (Fig. 3A lane 10), suggesting a shift in subunit composition towards H-rich heteropolymers (Fig. S5A). Indeed, the transcript levels of *Fth1* were markedly increased after LPS stimulation, whereas transcript levels of *Ftl* remained relatively unchanged (Fig. S5B). A similar increase in FTH1 levels was reported in monocytes exposed to inflammatory conditions.^30^ In agreement, the FTH1:FTL protein ratio shifted from approximately 1:1 to approximately 3:1 after LPS stimulation (Fig. S5C). In contrast, LPS stimulation of epithelial cells led only to a slight increase in *Fth1* transcript levels, alongside a significant increase in *Ftl* transcript levels (Fig. S5B). Nevertheless, epithelial ferritin protein concentration remained below detectable levels in immunoblots; only epithelial ferritin-iron could be detected, and this was decreased after 24 h of LPS stimulation (Fig. 3A). In summary, the complex dynamics of ferritin synthesis and degradation during inflammation match the increase in macrophage ferritin and ferritin iron, and the decrease in epithelial ferritin iron, and are in agreement with the observations in the inflamed lesion of *Tnf* ^*ΔARE/+*^ mice (Fig. 1A).

**Figure 3:**
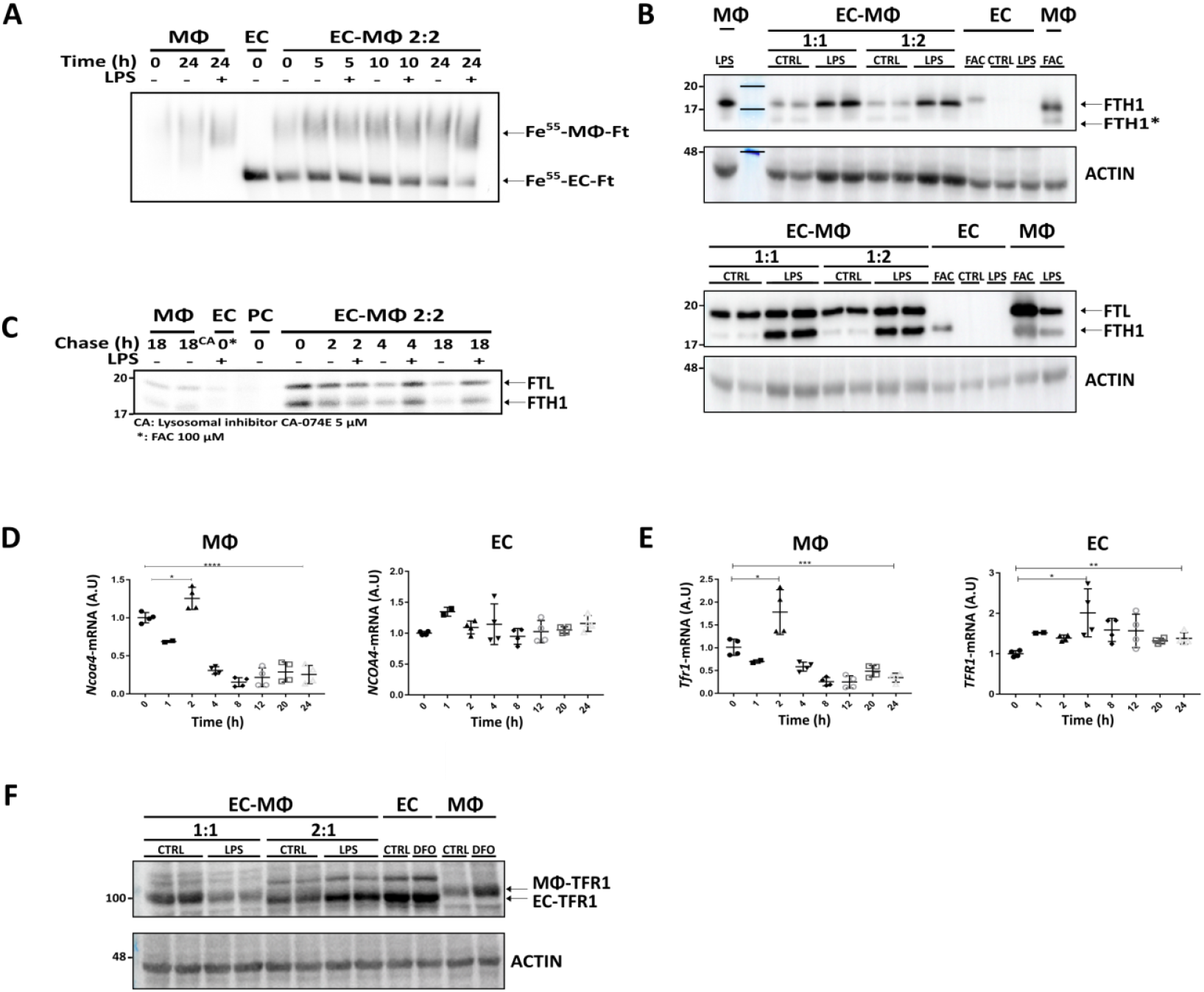
Iron-regulated proteins sense and respond to reduced epithelial and increased macrophage iron levels. Co-cultures of epithelial cells (EC) and macrophages (MΦ) were treated with 200 ng/ml lipopolysaccharide (LPS), ferric ammonium citrate (FAC), or desferrioxamine (DFO) for 24 h or as indicated. **(A)** Co-cultures were pretreated with ^55^Fe-labeled human transferrin for 16 h, and then washed and treated with LPS. Cell-lysates were separated on a native gel (n=2). A representative radiograph is shown. (**B)** Representative immunoblot for the ferritin H-subunit (FtH) and ferritin L-subunit (FtL). EC ferritin was not detected (except in FAC-treated ECs), thus the bands are MΦ ferritin only. Anti-FtL antibody detects both subunits (FtH is the lower band just above the 17kD Mw-marker) n=2. (**C)** Metabolic labeling (^35^S) pulse-chase experiment of LPS-stimulated EC-MΦ (BMDM) co-cultures, followed by ferritin immunoprecipitation (IP) (n=2). (**D)** RNA expression levels of *Ncoa4* were measured in co-cultures by RT-qPCR (n=4). **(E)** Transferrin receptor-1 (*Tfr1*) RNA expression levels were measured by RT-qPCR (n=4). (**F)** A representative immunoblot for TFR1 (n=12). In co-cultures, EC were always Caco2 cells. MΦs were as follows: BMDM in **A – E**, and Raw264.7 in **F**.

To further validate the observed inflammatory iron pattern in the epithelial-macrophage co-culture, changes in mRNA levels of the iron importer TFR1 were evaluated following LPS induction (Fig. 3E). We found that *Tfr1* mRNA and protein levels decreased in the macrophages and increased in the ECs 24 h after LPS exposure, consistent with the iron accumulation in the macrophages and the iron decrease in the IECs of the *Tnf* ^*ΔARE/+*^ mice. The *Tfr1* mRNA decrease in macrophages was preceded by an increase in *Tfr1* transcripts, identified 2 h after LPS stimulation, suggesting an early inflammation-mediated increase in iron import that contributes to iron accumulation in the macrophages (Fig. 3E-F). Taken together, in epithelial cells, the decrease in ferritin and increase in TFR1 are in line with the activation of IRP1. However, in the macrophages, the IRP1 activation was decoupled from high cellular iron status, and ferritin and TFR1 regulation were dominated by the iron-loaded phenotype.

### IRP1 activity contributes to the propagation of inflammation

Since IRP1 RNA-binding activity was increased in inflamed macrophages, regardless of their iron-loaded phenotype (Fig. 1-3), we hypothesized that this activation may play a role in the course of the inflammation. Thus, the effect of IRP1 knockout in macrophages on inflammatory markers and iron pattern was assessed. The classical inflammatory response of co-cultures involving *Irp1*^*-/-*^ macrophages was reminiscent of the co-cultures with WT macrophages in all inflammatory parameters tested, including the levels of secreted epithelial cytokine IL-8 and transcript levels of the macrophage inflammation markers *Tnfα, Icam1* and *Il6* (Fig.S6A and B). In parallel, NO levels in the medium of both co-cultures (EC with WT or *Irp1*^*-/-*^ macrophages) were similarly increased in response to LPS stimulation (Fig. S6C). Furthermore, FTH1 was significantly increased in macrophages of both genotypes in inflamed co-cultures (Fig.4A), while aconitase activities were significantly decreased in both co-cultures (Fig.4B), indicating that deletion of IRP1 in the macrophages alone is not responsible for the inflammatory changes in iron homeostasis.

**Figure 4:**
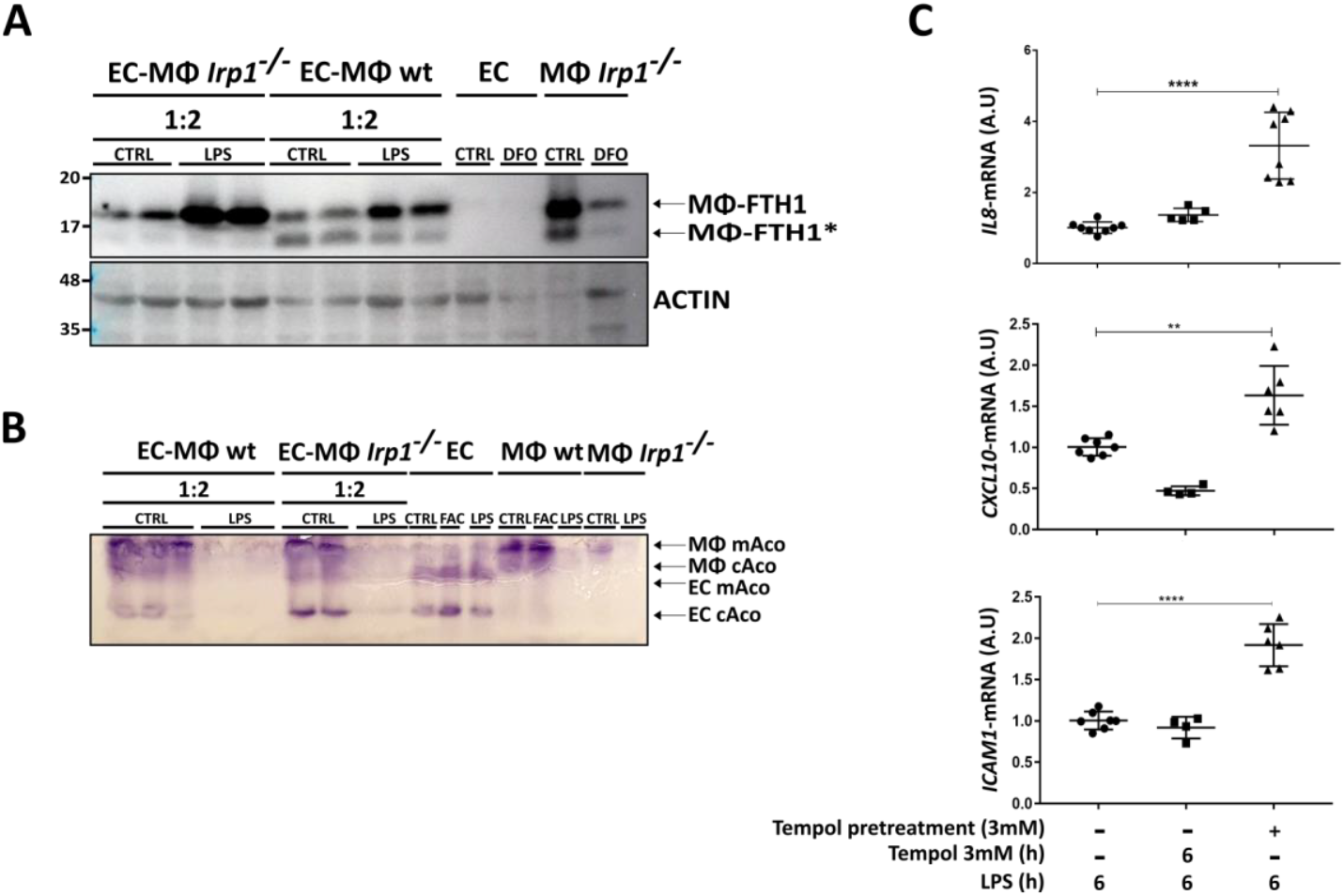
Macrophage and epithelial IRP1 IRE-binding plays different roles in intestinal inflammation. **(A) and (B)** Co-cultures of epithelial cells (EC) with WT-bone marrow derived macrophages (BMDM) or with BMDM with targeted deletion of IRP1 (MΦ *Irp1*^-/-^) were treated with 200 ng/ml lipopolysaccharide (LPS) and ECs and BMDMs were treated with ferric ammonium citrate (FAC), or desferrioxamine (DFO) for 24 h, as indicated **(A)** A representative immunoblot for FtH is shown (*FtH-degradation product) (n=8). **(B)** A representative gel of an **in**-gel aconitase assay is shown (n= 4). (**C)** ECs (Caco2) were pretreated with 3 mM Tempol for 16 h, washed, and then co-cultured with MΦs (Raw 264.7). After MΦs adhered, LPS and/or Tempol was added for 6 h. RNA expression levels were measured by RT-qPCR (n=6).

As epithelial cells have been shown to play a critical role in the pathophysiology of IBD,^31^ and epithelial iron deficiency has been associated with an inflammatory response,^32^ the effect of epithelial IRP1 activation on the inflammation was assessed. Activation of IRP1 was previously achieved by a stable aminoxyl radical Tempol.^33^ LPS stimulation of co-cultures with Tempol-pretreated epithelial cells, elicited a more significant inflammatory response in the epithelial cells compared to epithelial cells in co-cultures without Tempol pretreatment (Fig. 4C). Without LPS stimulation, Tempol treatment alone did not elicit a significant inflammatory response in the co-cultures. Yet, a trend of increased murine *Tnfα, Icam1* and human *ICAM1* was observed, when IRP1 was activated (Fig. S6D). These results suggested that epithelial IRP1 contributes to the course of the inflammation.

To test this, we introduced a targeted deletion of *Irp1* into the *Tnf* ^*ΔARE/+*^ mice. To easily monitor iron levels in the tissue sections, all mice were previously subjected to iron overload. As expected, a significant increase in LP ferric iron levels was observed in *Tnf* ^*ΔARE/+*^ mice compared to WT mice. In contrast, *Tnf* ^*ΔARE/+*^, *Irp1*^-/-^ mice showed no increase in LP iron levels similar to the non-inflamed LP in WT mice (Fig. 5A). In addition, *Tnf* ^*ΔARE/+*^, *Irp1*^-/-^ mice showed no histological signs of inflammation (Fig. 5B-C and S7A) and a significantly reduced pro-inflammatory response, as indicated by reduced *Tnfα* and *Icam1* levels (Fig. 5D). These results indicated that activation of IRP1 RNA-binding activity is required for the propagation of terminal ileum inflammation in *Tnf* ^*ΔARE/+*^ mice.

**Figure 5:**
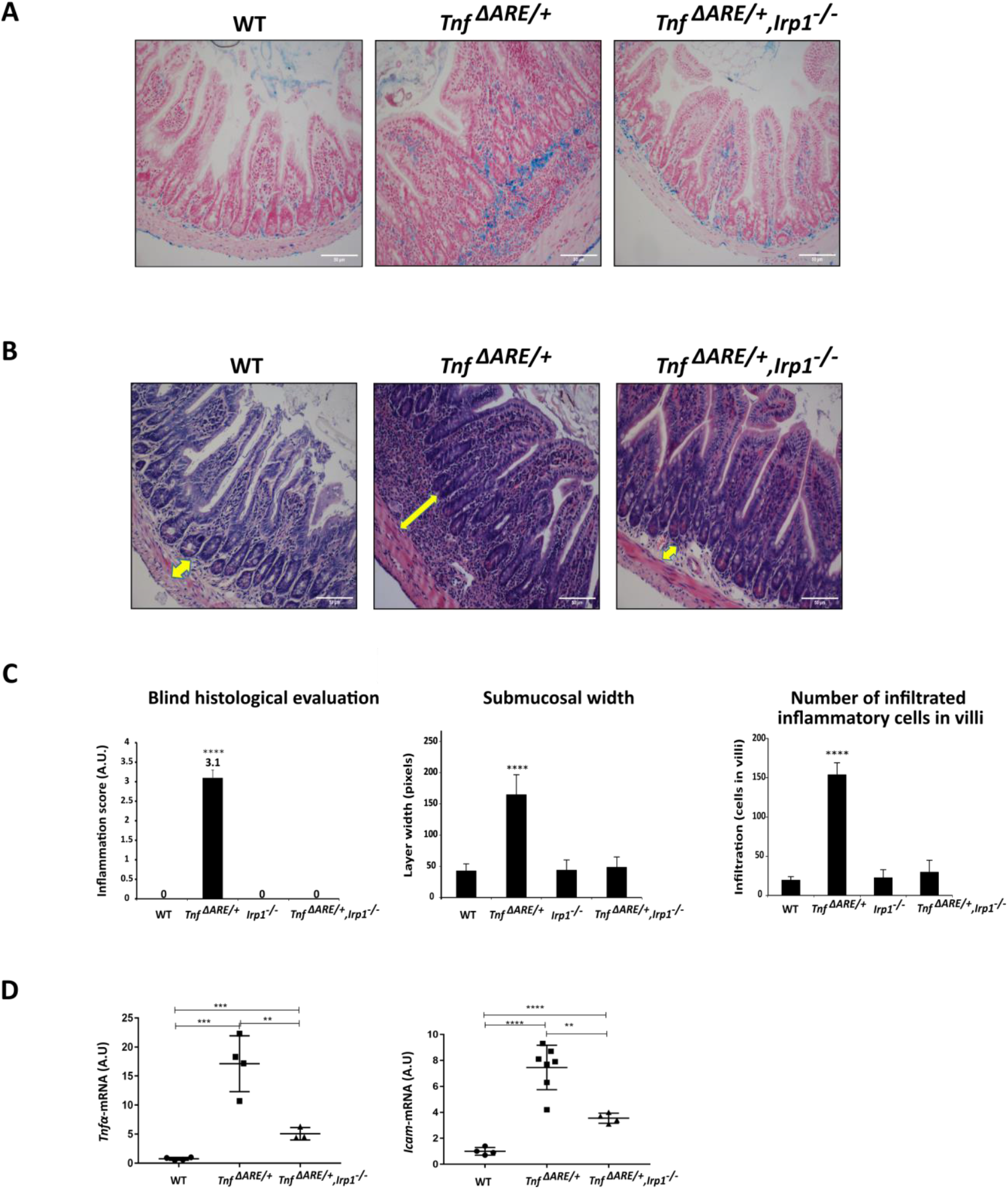
Deletion of IRP1 restores iron homeostasis and inhibits the inflammatory process in *Tnf ΔARE/+* mice. *Tnf* ^*ΔARE/+*^ mice were crossed with *Irp1*^-/-^ mice until *Tnf* ^*ΔARE/+*^, *Irp1*^-/-^ mice were created. Sections from terminal ileum (TI) of 12-14-week-old mice from indicated genotypes were prepared. **(A)** Representative images of Perls-stained TI sections of iron-overloaded mice, illustrating iron distribution *Tnf* ^*ΔARE/+*^, *Irp1*^-/-^ mice is similar to wild type and not to *Tnf* ^*ΔARE/+*^ mice. (**B)** Representative hematoxylin and eosin-stained TI sections illustrating normal histology in *Tnf* ^*ΔARE/+*^, *Irp1*^-/-^ mice, compared to *Tnf* ^*ΔARE/+*^ mice. The arrows highlight the width of the submucosal layer. **(C)** Blind histological evaluations of TI sections from 12-14 week-old WT, *Irp1*^-/-^, *Tnf* ^*ΔARE/+*^ and *Tnf* ^*ΔARE/+*^, *Irp1*^-/-^ mice (n= 6). (**D)** RNA expression levels of select proinflammatory cytokines were measured by RT-qPCR (n=4-7) in RNA from TI of 12-14 week-old WT, *Irp1*^-/-^, *Tnf* ^*ΔARE/+*^ and *Tnf* ^*ΔARE/+*^, *Irp1*^-/-^ mice.

### Modified iron homeostasis is observed in the inflamed lesions of CD patients

Finally, to determine whether the inflammatory iron homeostasis observed in the mouse models as well as in our co-culture models mimics the impaired iron homeostasis in CD patients, sections of intestinal biopsies from CD patients were labeled for ferritin. Indeed, ferritin levels proved to be significantly lower in IECs of inflamed compared to non-inflamed areas from the same patients (Fig. 6A, Fig. S7B). This suggested that also in IBD patients, iron homeostasis is mediated by inflammatory IRP1 activation. Thus, inhibiting IRP1 RNA-binding activity in inflamed intestinal lesions may result in a reduction in intestinal inflammation and serve as a novel approach to treat IBD.

**Figure 6:**
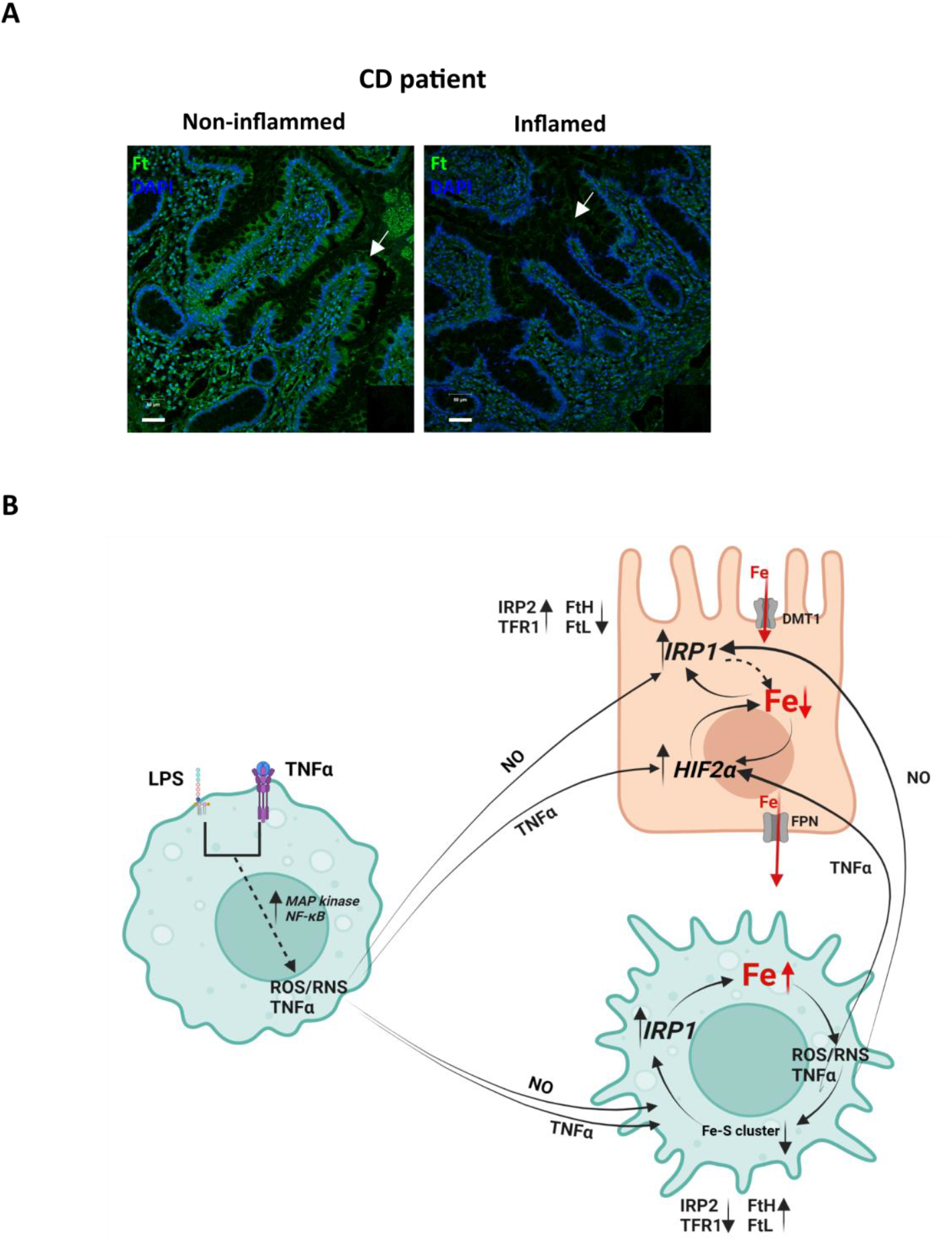
Modified iron homeostasis is observed in the inflamed lesions of Crohn’s disease patients. **(A)** Sections from non-inflamed and inflamed intestinal areas of Crohn’s disease patients (n=8) were immuno-stained for ferritin (green). Representative images show that the inflamed epithelial cells (IEC) (arrows) have decreased ferritin levels. Scale bars 50 μm. (**B)** A schematic presentation of the involvement of IRP1 in the self-perpetuating inflammatory process. Macrophages activated to a proinflammatory response secrete nitric oxide (NO) and proinflammatory cytokines, such as TNFα to their immediate environment, enabling them to reach neighboring cells, including epithelial cells. In epithelial cells, TNFα activates the hypoxia-inducible factor HIF2α, which mediates increased iron flux from the intestinal lumen to the lamina propria, while NO activates IRP1, which inhibits ferritin translation and thus inhibits ferritin storage, which further increases iron flux. In macrophages, NO activates IRP1 and thereby maintains a certain level of increased iron influx that continues to promote iron overload, which supports the pro-inflammatory polarization of the macrophage. The inflammation-mediated reduction in iron-sulfur cluster assembly in macrophages and increase in HIF2α in epithelial cells continue to support the activity of IRP1 and the iron flux from epithelial cells to macrophages, which maintains the inflammatory iron pattern and perpetuates inflammation.

## Discussion

Both, genetic and environmental influences play pivotal roles in the onset of IBD. However, the precise interplay through which these factors influence the composition of the gut microbiota, compromise the integrity of the intestinal barrier, recruit the immune system and trigger the inflammatory process, remains elusive. Nonetheless, it has been established that local immune cells undergo activation, leading to the release of NO, alongside an array of other inflammatory molecules and cytokines in the affected region during the initial stages.^34^ Here we demonstrated an inflammation-mediated activation of IRP1 in both, IEC and LP, driven by NO and decoupled from the cellular iron level in macrophages. We further observed that IRP1 activation led to an impaired iron homeostasis in IBD.

The pathological iron distribution was observed in human biopsies from CD patients, and in mouse models for CD and colitis and was replicable in cellular co-cultures. It is characterized by decreased iron stores in IECs and increased iron stores in LP macrophages. Inflammation is a response to structural or functional insults, and an integral part of animal physiology, particularly as a means to correct homeostatic instability.^35^ The iron shift induced a homeostatic instability that eventually perpetuated an inflammatory process, as both increased and decreased iron levels were shown to mediate inflammation.^3–5^

The data suggest that the inflammation-associated iron pattern was driven by the activation of IRP1. Similarly, in a model of non-alcoholic fatty liver disease, elevated IRP1 RNA-binding activity was found despite high iron levels in rat livers,^36^ demonstrating activation of IRP1 in an inflammatory context, decoupled from iron status. In IBD, increased ROS/RNS release from activated macrophages and reduced Fe-S cluster assembly can explain the IRP1 activation.^37^ The presented results demonstrated iron-decoupled activation of IRP1 in macrophages, and suggested that this activation signals a pseudo-iron deficiency, contributing to further iron influx.^6^

In epithelial cells, HIF2α was activated (Fig 2D), in line with reports of NFkB-p65 activation, eliciting the activation of HIF2α.^38^ HIF2α activation is possibly further supported by NO-mediated iron efflux,^39^ which leads to prolyl-hydroxylase deactivation and HIF2α stabilization.^40^ HIF2α activation induces increased iron flux, which includes iron import through the apical membrane and iron export from the basolateral membrane, securing the repletion of body iron levels in iron deficiency.^28^ This increased iron flux may further contribute to the reduced iron levels in the epithelial cells and the ensuing activation of IRP1. In addition, drug-induced activation of IRP1 in epithelial cells increased the production of proinflammatory cytokines by these cells (Fig. 4D). Thus, the activation of IRP1 RNA-binding activity plays a role in the establishment of an inflammatory iron-homeostasis that supports the exacerbation of the inflammatory process, as summarized in Figure 6B. This model is supported by our key finding that deletion of IRP1 abolished inflammation in *Tnfα*-overexpressing mice (Fig. 5A-D). Inflammation-driven regulation of iron homeostasis is thus a factor that plays an active role in the series of events that contribute to the course of inflammation.

Direct interference with intestinal iron homeostasis, by administering a low-iron diet, reduced inflammation in *Tnf* ^*ΔARE/+*^ mice.^41^ Interestingly, systemic iron chelation did not have a similar effect,^41^ supporting a role for local iron homeostasis in the course of the inflammation. Further, decreased iron content in the intestinal lumen elicited a lactobacillus-mediated reduction of iron uptake by decreasing HIF2α activity.^42^ Overall, this suggest a connection between the microbial composition or activity with epithelial iron homeostasis and thus also epithelial IRP1 activity. In addition, local regulation of iron homeostasis was also demonstrated to play an important role in the pathophysiology of IBD. For example, local production of hepcidin, (generally regarded as a systemic iron regulator) by conventional dendritic cells in the colon, regulates luminal iron content and supports colonic tissue repair.^43^

The presented data demonstrating opposing inflammatory regulation of iron homeostasis in macrophages versus epithelial cells in the same tissue, complicated studying mechanisms that drive iron redistribution. We therefore looked for a cellular co-culture system featuring the same inflammation-associated iron distribution pattern. Many of the published co-culture setups mimicked well-established inflammatory parameters, however did not replicate the inflammatory iron pattern. This work showed that the inflammatory iron homeostasis was only established when epithelial cells and macrophages were in direct contact. This cell-to-cell contact enabled us to replicate the iron and ferritin distribution that was observed in the *Tnf* ^*ΔARE/+*^ and DSS mouse models and IBD patients. Similar inter-species co-cultures of Caco2 cells and Raw264.7 cells were successfully applied to analyze inflammatory pathways.^22^ Also, human grafts of fetal intestine in SCID mice were used to study fistulae in IBD,^44^ supporting the functionality of human-mouse interspecies systems. Interspecies co-cultures have many advantages, especially in our study, they enabled us to differentially analyze iron-related proteins from epithelial versus macrophage origin. Using this model, we found that in response to the induction of inflammation, TFR1 was regulated by the iron status of the specific cell types (elevated in ECs and decreased in macrophages), as seen in the *Tnf* ^*ΔARE/+*^ animal model. The interspecies co-culture also enabled analysis of inflammation-mediated changes in ferritin. In the macrophages, significant increases in both ferritin protein and ferritin-iron, as well as a shift in the ferritin composition towards FTH1-rich complexes were measured in response to inflammation. Considering that NCOA4 binds specifically to FTH1, and not to FTL,^45^ the change in subunit composition may have important implications on ferritin trafficking. In light of the recent findings that ferritin-NCOA4 complexes generate liquid-phase condensates,^46,47^ and taking into account that ferritin is an abundant and large cytosolic protein, the inflammation-specific ferritin subunit ratio may alter the physicochemical properties and the mesoscale dynamics of the cytosol in inflammation.

In the physiologic, non-inflamed state, IRP2 plays a dominant role in iron regulation, and deletion of IRP1 plays only a transient role.^12,14,48,49^ In addition, reduction of IRP1 protein levels and activity in the non-inflammatory setting of superoxide dismutase 1 knock-out mice was previously reported to have no effect on iron homeostasis.^50^ Yet, in patients with genetic hemochromatosis, dysregulation of IRP1 and IRP2 has been observed and was attributed to the release of NO produced in activated macrophages,^17^ and in our hands, when activated by inflammatory signals, IRP1 acts as a dominant regulator of iron homeostasis of inflammation.

## Conclusions

Taken together, the presented findings demonstrated that an inflammation-induced shift in the regulation of cellular iron homeostasis is mediated by the activation of IRP1, and that this shift plays an active role in the propagation of inflammation in IBD. Further, correction of iron homeostasis by interfering with IRP1 activity halted the self-perpetuation of inflammation. Thus, unlike conventional therapeutic approaches which focus on modulating the immune system, our findings may pave the way for new therapeutic interventions for IBD by targeting IRP1.

## Acknowledgements

We like to thank Dr. T.A. Rouault (NICHD, NIH, Bethesda MD, USA) for providing the IRP1^-/-^ mice, Dr. F. Cominelli (Case Western Reserve University, Cleveland, OH, USA) for initially providing the *Tnf* ^*ΔARE/+*^ mice, and Dr. G. Kollias (BSRC Alexander Fleming Athens University Medical School, Athens, Greece) for subsequently sharing with us the *Tnf* ^*ΔARE/+*^ mice. Drs. Dan Gelvan, Vasiliki Koliaraki, Benjamin Podbilewicz and Dan Cassel and many former and present lab members of the E.M-H. lab for critically reviewing the manuscript and supporting this project with their thoughts and discussions.

## Author contributions

Conceptualization: EGMH

Methodology: LF, SB, WZ, NGR, NG, AN, AZ, MW, RW

Investigation: LF, SB, WZ, WHT, KL, NGR, NG, AN, AZ, MW, RW

Visualization: LF, SB, WZ, AZ

Funding acquisition: EGMH, NGR

Project administration: EGMH, LF, SB

Supervision: EGMH

Writing – original draft: LF, SB, WHT, EGMH

Writing – review & editing: EGMH, LF, WHT, KL, NGR, NG

## Data and materials availability

All materials are available from the authors on reasonable request with materials transfer agreements (MTAs).

## Supplementary Materials

Materials and Methods

Figs. S1 to S7

Tables S1 to S5

References (*66–70*)

