## Supplementary data for "Iron Regulatory Protein 1 is Required for the Propagation of Inflammation in Inflammatory Bowel Disease"

**The PDF file includes:**

Figs. S1 to S7
Materials and Methods
Tables S1 to S5
References (1-5)

### Supplementary Figure 1

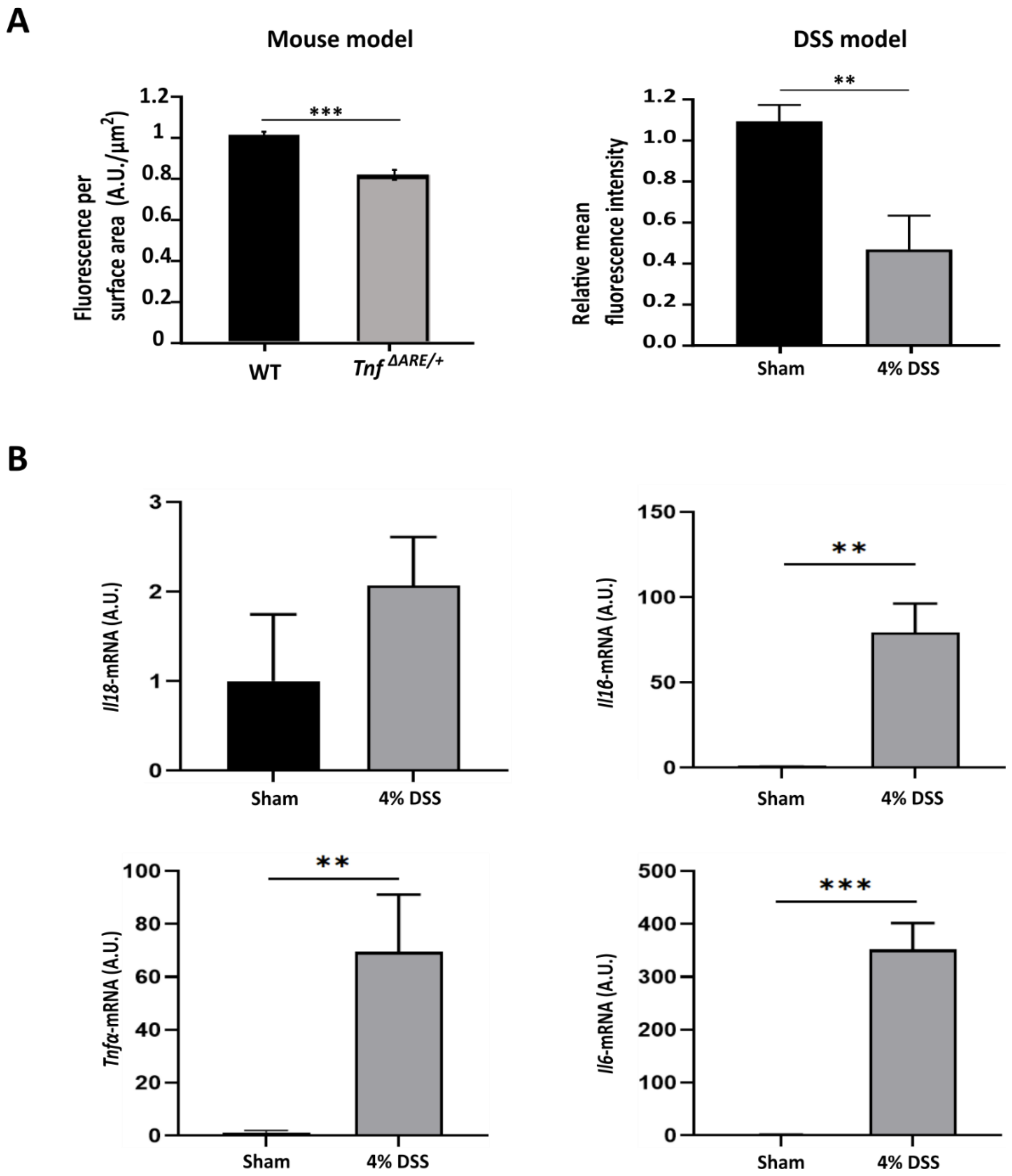


**Supplementary Figure 1: Ferritin levels in inflamed epithelium.**
**(A)** To further corroborate the measured decrease in ferritin levels in the epithelial layer, the fluorescence intensity of epithelial ferritin (Ft) was quantified in immunofluorescent stained sections (Fig. 1A) using IMARIS (sections from *Tnf* $^{\Delta ARE/+}$ mice and WT controls) and Image J (DSS and sham mice) software. Fluorescence of the inflamed epithelium is presented as a fraction of the Ft intensity measured in control epithelium (mean ± SEM (n=3-4/group)). (**B**) RNA expression levels of select proinflammatory cytokines were measured by RT-qPCR (n=3/group) and were significantly increased in the colon of DSS- compared to sham-treated mice.

### Supplementary figure 2

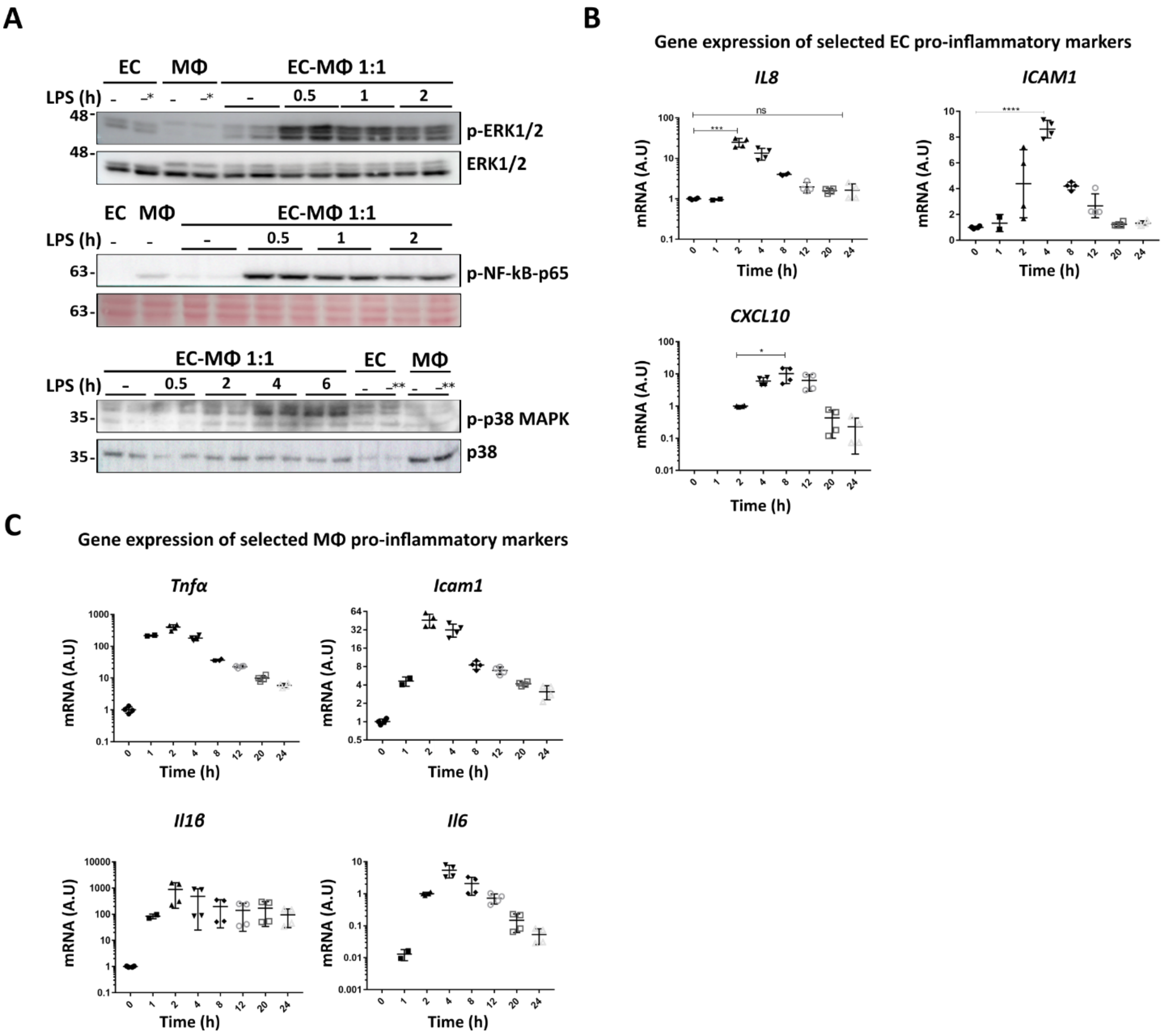

**Supplementary Figure 2: Validation of the inflammatory response in co-cultures of epithelial cells (EC) with macrophages (MΦ).**

The human epithelial colon cancer line, Caco-2 cells, were co-cultured with either the mouse macrophage cell line Raw 264.7 or with bone marrow-derived primary macrophages (BMDM). To distinguish between data from tissue fractions (isolated from mouse intestine) and data from co-cultures, we denote intestine-derived fractions enriched with intestinal epithelial cells as IECs and and refer to the epithelial Caco-2 cell-line as EC. The intestine-derived fraction enriched with immune cells from the lamina propria is referred to as LP. Both sources of macrophage are referred to as MΦ. **A-C** Co-cultures of EC and MΦ were treated with 200 ng/ml LPS for the indicated time intervals. **(A)** Representative immunoblots of ERK 1/2, p-p38 MAPK, p-38 MAPK, and p-NFkB-p-65, using lysates from co-cultures of Caco-2 cells with BMDM and their corresponding mono-cultures. LPS stimulation caused a rapid increase (peak within 30 min) in p-ERK1/2 and p-p-65 (NFkB) levels. Phosphorylation of p-38 peaked 4 h after LPS stimulation. (**B-C)** RNA expression levels of select pro-inflammatory cytokines were measured in Caco-2-BMDM (MΦ) co-cultures by RT-qPCR (n=4). LPS stimulation led to increased expression of *IL8*, *ICAM1*, and *CXCL10* in EC (Caco-2 cells) and *Tnfα*, *Icam1*, *IL1b* and *Il6* in macrophages (BMDM). All primers were tested for species-specificity and no cross-reaction between primers for human or mouse genes was detected. *Cells were cultured in 21% oxygen. All others were cultured in 6% oxygen. **Cells were treated with 100 μM DFO for 24 h. In co-cultures, EC were always Caco2 cells. MΦs were as follows: Raw264.7 in **A** (ERK and NFkB-p65 gels) and BMDM in **A** (p38 gel MAPK) and **B** and **C**.

### Supplementary figure 3

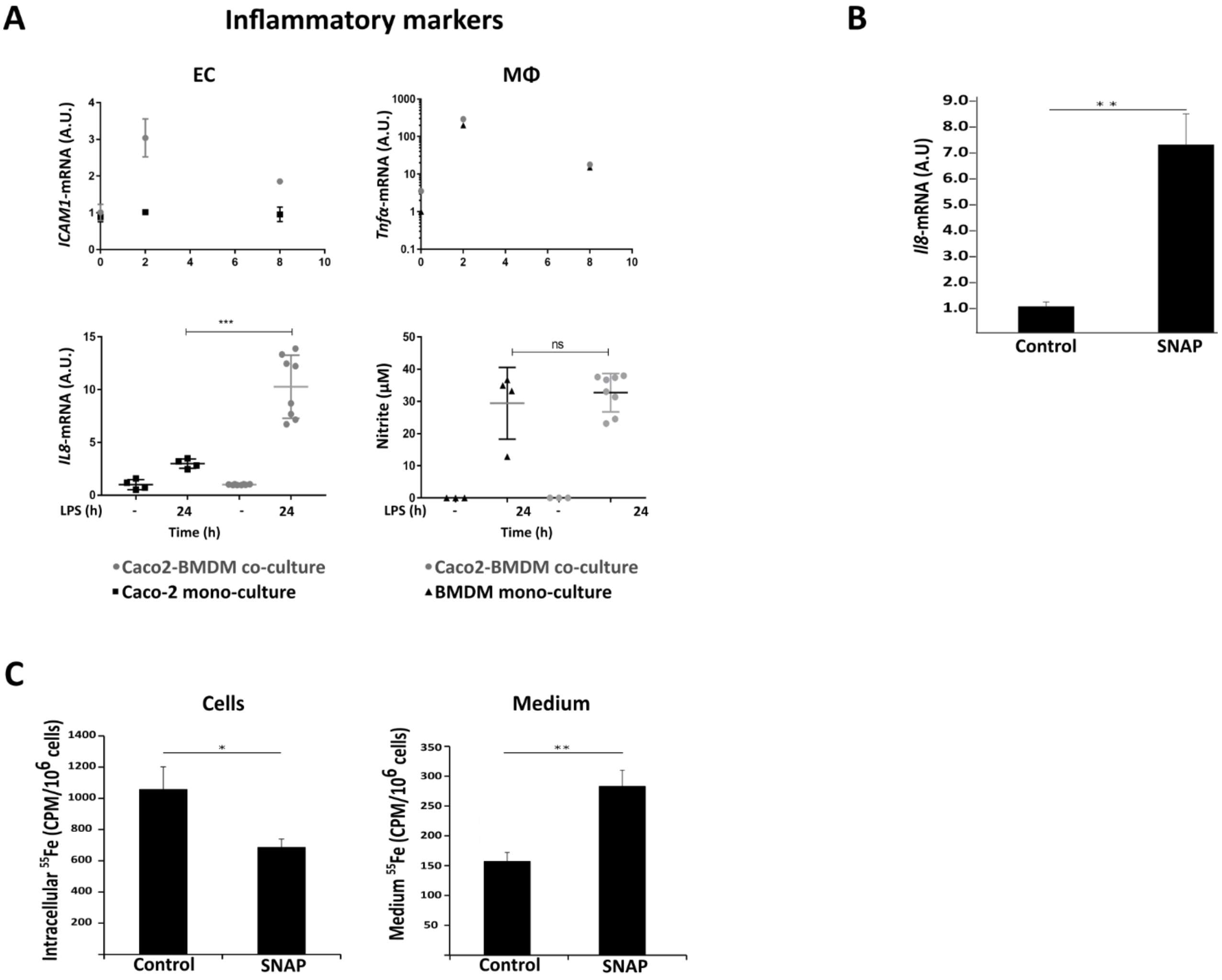


**Supplementary Figure 3: Iron phenotype in epithelial inflammation:**
As LPS easily triggers macrophages but not Caco-2 cells to produce proinflammatory cytokines, we tested whether the inter-species co-culture can activate Caco-2 cells. **(A)** Co-cultures of EC (Caco2) with MΦ (BMDM) and their corresponding monocultures were treated with LPS for the indicated time intervals. RNA expression of select genes was measured by RT-qPCR. In EC monocultures, LPS-triggered activation of genes was not identified, yet was identified in both MΦs monocultures and in MΦ-EC co-cultures. Further, IL-8 that is secreted by activated ECs, was only found in the medium of the co-cultures. Nitric oxide was produced by LPS-activated MΦs and diffuses to the medium and to neighboring cells in close vicinity. Nitrite, an oxidation product of nitric oxide was detected in MΦ mono-cultures and in co-cultures 24 h after LPS treatment. **B-C.** ECs were treated with a nitric oxide donor, S-nitrosoN-acetylpenicillamine (SNAP) for 22 h in two sequential doses. **(B)** *IL8* RNA expression was measured by RT-qPCR. Expression levels were elevated in ECs after SNAP treatment (mean ± SD (n=3)). (**C**) Intracellular (left) and extracellular (right) $^{55}Fe$ after SNAP treatment of ECs. $^{55}Fe$ levels were measured by a beta-scintillation counter (mean ± SD (n=6)). SNAP induced a decrease in intracellular iron levels by promoting iron secretion. These data suggest that nitric oxide either directly or indirectly (such as via activated IRP1) causes epithelial iron depletion and increases iron secretion from ECs.

### Supplementary figure 4

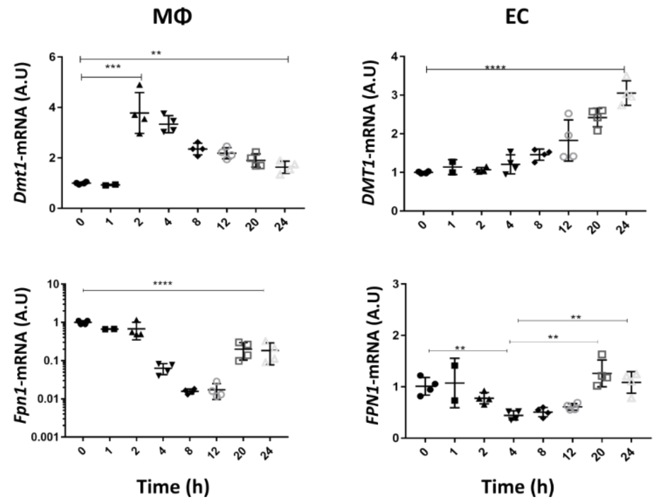


**Supplementary Figure 4: Inflammation mediates transcriptional regulation of the iron importer divalent metal transporter (DMT)1 and the iron exporter ferroportin (FPN)1.** Co-cultures of epithelial Caco-2 cells (EC) with BMDM macrophages (MΦ) were treated with 200 ng/ml LPS for the indicated time intervals. Expression of the iron importer *Dmt1* and iron exporter *Fpn1* RNA transcripts were measured by RT-qPCR (n=4). Expression levels of *Dmt1* in MΦs were strongly increased 2 h after LPS stimulation and slowly decreased during the subsequent 24 h, still remaining at a significant 2-fold increase. The regulation pattern of *Dmt1* was reminiscent of that of *Tfr1* (Fig. 3E) in MΦs, both supporting an initial increase of iron import through the TFR1/DMT1 pathway, initiated by IRP1 activation and transcript stabilization. In ECs, *DMT1* and *FPN1* were strongly regulated by HIF2α, and both genes were significantly induced, starting at 4 h, when HIF2α activation was detectable (Fig. 2D). As FPN1 is strongly regulated at many posttranscriptional levels, its strong decrease in MΦs (note the logarithmic scale of the y-axis in this graph) supports the iron retention in these cells, while in epithelial cells, the recovery from an initial decrease supports continuous epithelial iron export through FPN1.

### Supplementary figure 5

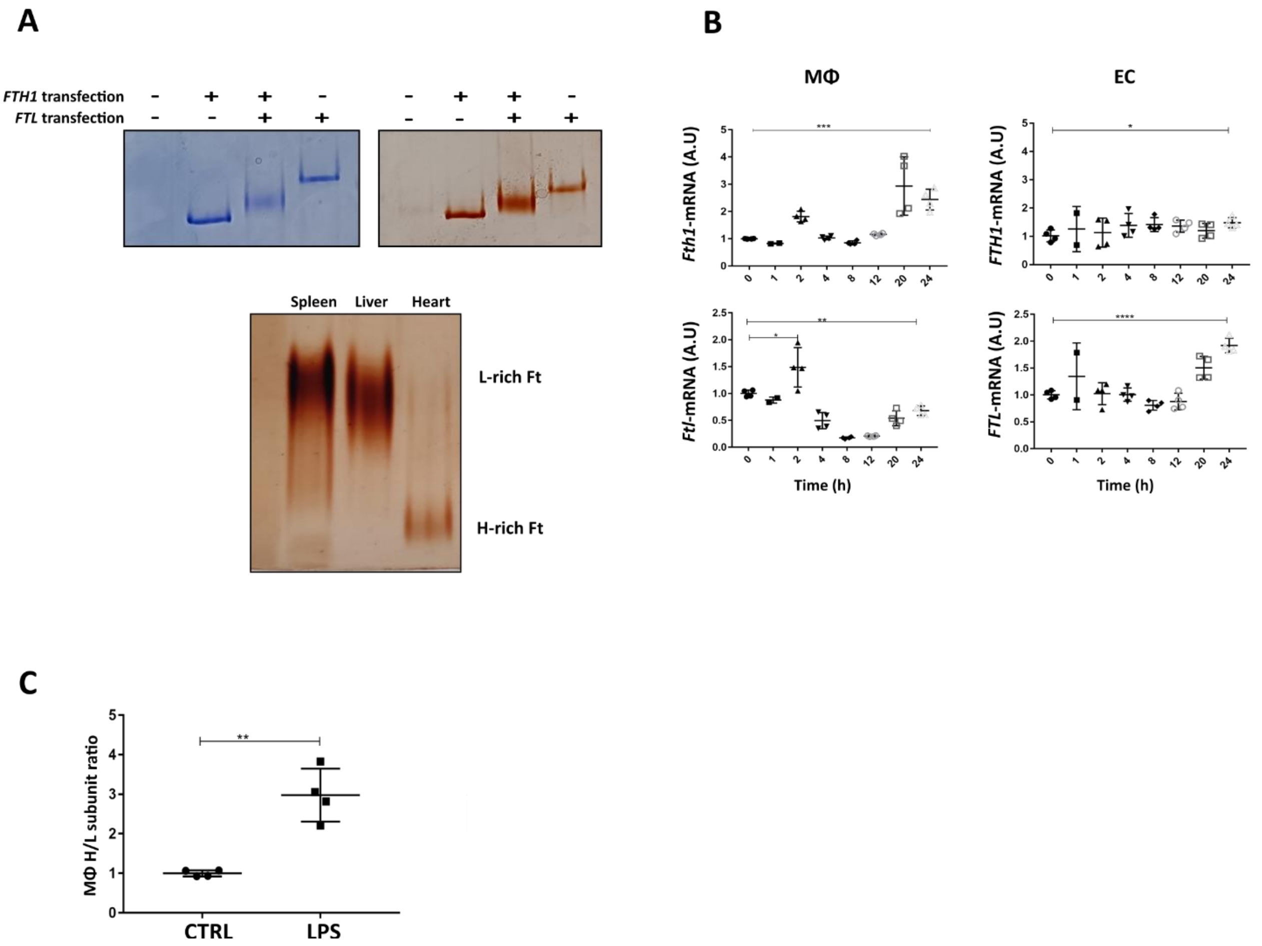


**Supplementary Figure 5: Ferritin subunit composition in inflammation.**

The strong increase in FTH1 and the slight increase in FTL in response to LPS stimulation led us to further investigate the subunit composition of ferritin. **(A)** Technical ability to visualize ferritin complexes with different ferritin subunit compositions: Cell lysates (upper panel) and lysed tissues harvested from C57BL/6 mice (lower panel) were subjected to heat treatment to partially purify ferritin before analysis on native gels. Upper panel: HEK293T cells were transfected with the indicated constructs. Coomassie blue stain (left) and Perls-DAB stain (right). Endogenous ferritin was below detection levels in HEK293T cells (left lane). FtH:FtL (1:1) appears as a smear, suggesting assembly to diverse subunit combinations. Lower panel: Perls-DAB stain of tissue lysates demonstrate how ferritins rich in L-subunit migrate slower than ferritins rich in H-subunit, due to the differences in the isoelectric points of the two subunits. The slight shift of macrophage ferritin, 24 h after LPS stimulation is visible in Fig. 3A and suggests a shift in the subunit composition of macrophage ferritin towards a more FTH1-rich complex, supporting the calculation shown in Fig. S5C. (**B)** Co-cultures of Caco-2 epithelial cells (EC) with BMDM macrophages (MΦ) were treated with LPS 200 ng/ml for the indicated time intervals. RNA expression levels were measured by RT-qPCR (n=4). Transcript levels of *Fth1* increased significantly in LPS-stimulated MΦs, and only slightly in the ECs. In contrast, *Ftl* transcripts decreased in the MΦs and strongly increased in ECs. (**C)** Protein quantification of Fth1 and Ftl levels were based on the immunoblots and analyzed by ImageJ (n=4). Considering that at baseline, the subunit ratio is approximately 1:1 (as seen in the metabolic labeling in Fig. 3C, lane 5), we calculated the relative change of the subunits, comparing to the subunit in the non-stimulated state. The ratio between Fth1 and Ftl subunits in the ferritin complex was three times higher after LPS stimulation compared to control.

### Supplementary figure 6

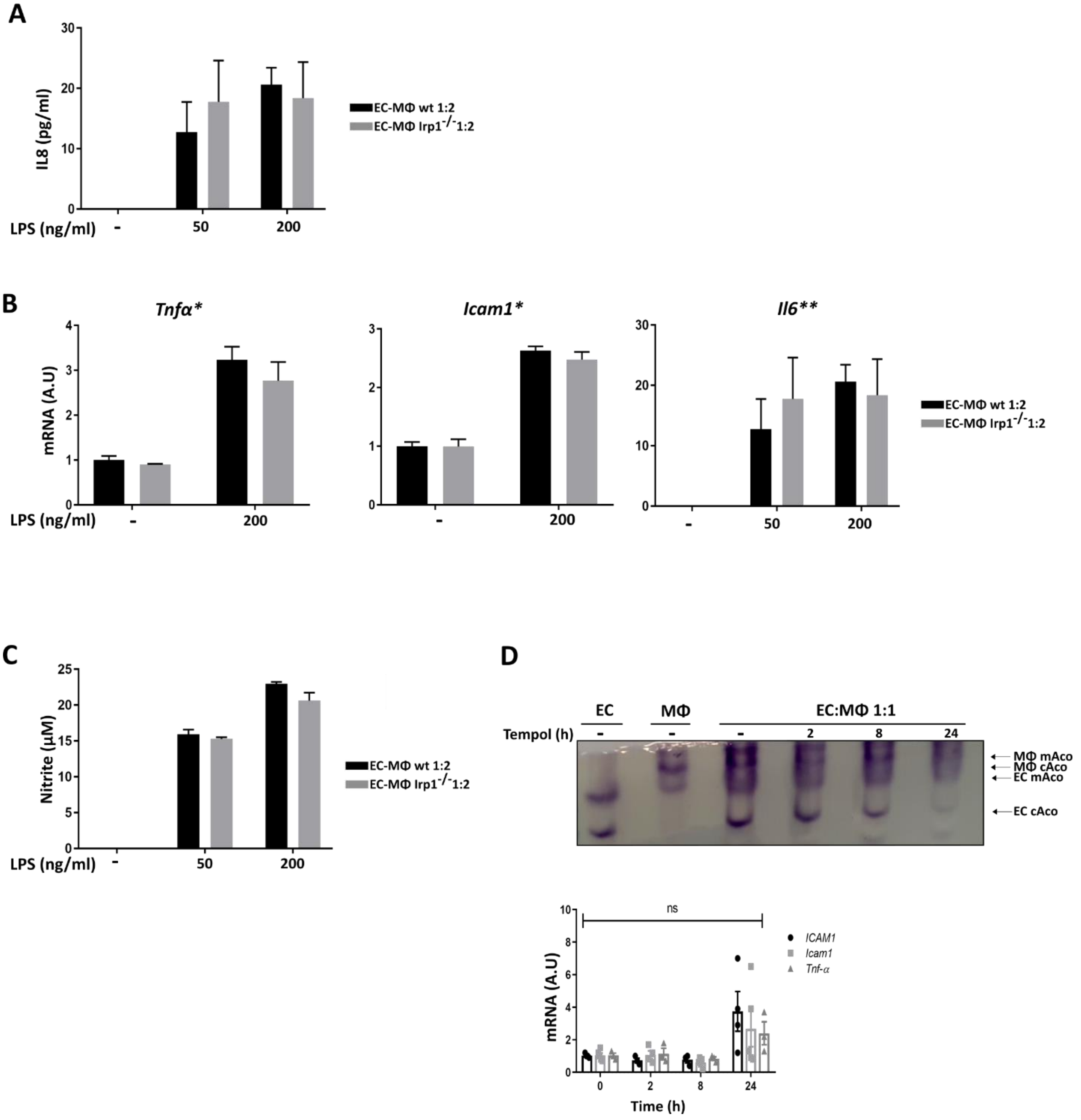

**Supplementary Figure 6: Manipulation of IRP1 activity.**

As IRP1 RNA-binding activity was activated in the macrophages, despite iron accumulation in these cells, we first hypothesized, that targeted deletion of IRP1 in these cells may be sufficient to manipulate the inflammatory iron distribution pattern and with it, the inflammation. **A-C.** Co-cultures of Caco2 (EC) with BMDM (MΦ) from WT mice and of EC with MΦ from *Irp1*$^{-/-}$ mice were treated with the indicated concentrations of LPS. (**A)** IL-8 levels were measured by ELISA (n=6). The increase in IL-8 levels after LPS stimulation was not significantly different in EC- WT MΦ compared to EC-*Irp1*$^{-/-}$MΦ co-cultures. (**B)** RNA expression levels of select pro-inflammatory cytokines were measured by RT-qPCR (n=2) and found to be similar in EC-wtMΦ compared to EC-*Irp1*$^{-/-}$MΦ co-cultures. *Expression calculated relative to the control WT co-culture. **Expression calculated relative to WT co-culture stimulated with 50 ng/ml LPS. (**C)** Nitrite concentrations in the medium were assessed using the Griess assay (n=6). Nitrite concentrations after LPS stimulation were not significantly different in EC- WT MΦ compared to EC-*Irp1*$^{-/-}$MΦ co-cultures. (**D)** EC-MΦ (Raw 264.7) co-cultures were treated with Tempol (3 mM) for the indicated time intervals. The representative gel shows that following Tempol treatment, while there was no decrease of aconitase activity in the MΦs, there was a significant decrease in ECs. This suggested that MΦs were not sensitive to the concentration of Tempol applied. RNA was isolated from Tempol-treated co-cultures and RNA expression of select pro-inflammatory cytokines (*Tnfα* and *Icam1* from MΦs and ICAM1 from ECs) was evaluated by RT-qPCR.

### Supplementary figure 7

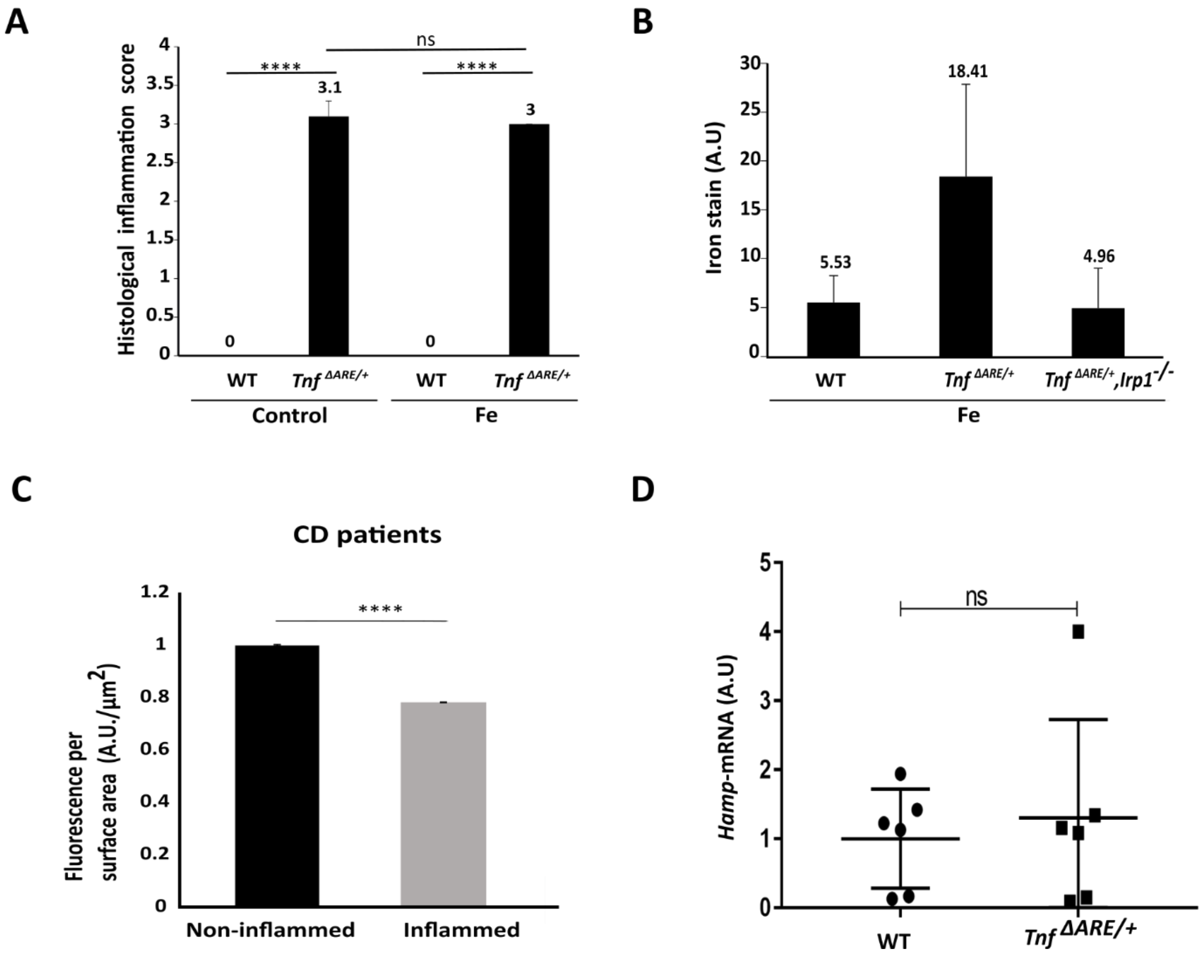


**Supplementary Figure 7: Deletion of IRP1 completely abolishes inflammation in *Tnf* $^{\Delta ARE/+}$mice.**

**(A)** Evaluation of sections from *Tnf* $^{\Delta ARE/+}$mice, with and without iron overload, was performed by a blinded board-certified pathologist and inflammation was scored according to standard histologic criteria, as previously described (*4*). Iron overload did not exacerbate the inflammatory phenotype of *Tnf* $^{\Delta ARE/+}$ mice. n=7 for WT; n=5 for *Tnf* $^{\Delta ARE/+}$ and WT-Fe; n=4 for *Tnf* $^{\Delta ARE/+}$-Fe. No statistical difference was found between control and iron overloaded (Fe) mice. (**B)** Quantification of mice shown in Fig. 5A. Perls-stained iron in terminal ileum sections of iron-overloaded mice (n=8). (**C)** The fluorescence intensity of epithelial ferritin (Ft) in the stained sections was evaluated using IMARIS software, and is presented as the ratio to Ft intensities in non-inflamed sections of the same CD patient. The ratio is presented as mean ± SEM. (n=8), (**D)** Liver hepcidin levels were evaluated by RT-qPCR. Values are presented as mean ± SD (n=6). Liver hepcidin mRNA levels were not significantly different in *Tnf* $^{\Delta ARE/+}$ (inflamed) compared to WT mice.

#### Supplemental Materials and Methods

##### Mice

Housing: Mice were housed in specific pathogen-free animal facilities at the animal facility of the Technion, Haifa, Israel, and studies were carried out in accordance with the recommendations in the Guide for the Care and Use of Laboratory Animals of the pre-clinical research authority at the Technion, Haifa, Israel. The protocols were approved by the Technion Animal Ethics Committee, Haifa, Israel (IL-135-09-19). For the dextran sulfate sodium (DSS) model, the protocols were approved by the Animal Investigation Ethics Committee of the affiliated Drum Tower Hospital of Nanjing University Medical School and were performed according to the Guidelines for the Care and Use of Laboratory Animals published by the National Institutes of Health, USA.

Mice with genetic modifications: IRP1 knockout ($Irp1^{-/-}$) mice on C57BL/6 background ($Irp1^{-/-}$) were a generous gift from Dr. Tracey Rouault (Molecular Medicine Program, National Institute of Child Health and Human Development, NIH, Bethesda, MD USA). $Tnf^{\Delta ARE/+}$ mice on C57BL/6 background ($Tnf^{\Delta ARE/+}$) were received initially from Prof. F. Cominelli at Case Western University and later from Prof. George Kollias, BSRC Alexander Fleming Athens University Medical School. $Tnf^{\Delta ARE/+}$ mice with targeted deletion of *Irp1* were generated by crossing $Tnf^{\Delta ARE/+}$ mice with $Irp1^{-/-}$ mice. Wild type (WT) littermates from breeders of $Tnf^{\Delta ARE/+}$ with WT mice and age and sex matched $Irp1^{-/-}$ mice were used as controls in the experiments described here.

Systemic iron overload of mice: Fe-dextran (Sigma) (90 mg/ml) was injected intra-peritoneally, daily for five days into 12-13-week-old mice as previously described [1]. The mice were sacrificed three days after the last injection.

DSS model: DSS with average molecular weight of 36,000-50,000 Da (MP Biomedicals) was added for seven days to the sterile drinking water of 7-10-week-old C57BL/6 (WT) male mice at a concentration of 4% and was administered *ad libitum*. Body weight and blood in stool were monitored daily.

##### Tissue collection and handling

Mice were euthanized and tissues (terminal ileum (TI) and distal colon) were collected and gently rinsed with cold Phosphate-buffered saline (PBS) solution. Samples were either fixed in 4% paraformaldehyde (PFA, Barnaor) for histological experiments, or quickly frozen in liquid nitrogen and then stored at -80 ℃ for RNA and protein extraction.

##### Tissue fixation, immunohistochemistry, and immunofluorescence staining

Mouse tissues were fixed in 4% PFA solution for at least 16 h, then dehydrated with 70% ethanol. Tissues were subsequently embedded in paraffin by the Rambam Medical Center Pathology Unit, Haifa, Israel. Blocks were then sectioned (5 µm-thick) using a Shandon Finesse 325 Manual Rotary Microtome (Thermo Fisher Scientific). Sections were fixed onto Superfrost microscope slides and later deparaffinized with xylene and rehydrated in a series of graded ethanol. For the DSS experiments, tissues were handled similarly, and then cut to 3 µm-thick sections.

##### Mouse tissue histology

###### Hematoxylin and eosin (H&E) staining

For H&E staining, paraffin-embedded terminal ileum sections were deparaffinized with xylene and rehydrated in 100% ethanol and then in 95% ethanol. Staining was performed with Harris hematoxylin solution (Sigma) and eosin Y (Sigma), as per the manufacturer’s instructions.

**Perls' Prussian blue iron staining**

Paraffin-embedded TI sections prepared as described, were incubated with freshly prepared Perls' Prussian blue solution (2% hydrochloric acid mixed with 2% potassium ferrocyanide) to reveal iron content and distribution. Quantification was evaluated by morphometric means as described previously [2].

**Assessment of inflammatory bowel disease (IBD)**

H&E-stained sections were graded (0: No change; 1: Minimal; 2: Mild; 3: Moderate; 4: Severe) by a blinded board-certified pathologist, based on standard histologic criteria [3].

**Mouse tissue immunofluorescence (IF)**

**Mice with genetic modifications:** Paraffin-embedded TI sections, prepared as described above for the iron stain, underwent antigen retrieval by incubation in citrate buffer (1x) in a microwave for approximately 10 min until boiling was observed. Following this, the sections were thoroughly rinsed with distilled water (dd$H_2O$) and blocked with 10% normal goat serum (Jackson) in PBS containing 0.1% bovine serum albumin (BSA, Sigma). For immunofluorescent staining, the sections were incubated overnight at room temperature (RT) in a humidified chamber with an affinity-purified rabbit anti-mouse liver ferritin antibody (1:200 dilution), kindly provided by Prof. A. M. Konijn. Afterward, the slides were washed three times with 1 x PBS. To detect the primary antibody, Alexa Fluor® 488-conjugated anti-rabbit secondary antibodies (A11034, Invitrogen) were used at a dilution of 1:1000 in PBS containing 0.1% BSA. The secondary antibodies were incubated with the sections for 1 h at RT. Following the incubation, the slides were washed three times with 1 x PBS for 5 min. The stained sections were mounted with VECTASHIELD mounting medium containing DAPI (4′,6-diamidino-2-phenylindole, VE-H-1200, Vector laboratories) to visualize the nuclei. Negative controls were included, where sections were incubated only with secondary antibodies. Microscopic imaging and fluorescent visualization were conducted using a Nikon Eclipse 50i microscope equipped with an X-cite series 120 microscope light source system. The acquisition software used for imaging was NIS-Elements Microscope Imaging Software. The fluorescent signal was quantified using IMARIS software.

**DSS model**: IF staining of ferritin in DSS treated mice and their controls, tissue sections (3 μm-thick) were utilized. The tissue sections were incubated overnight at 4 °C with a rabbit anti-ferritin heavy chain (FtH) antibody, which was in-house produced at Kuanyu Li's lab, Nanjing University Medical School. The primary antibody was diluted 1:1000 in PBS containing 1% BSA. Following the incubation with the primary antibody, the slides were incubated with Alexa Fluor® 488 AffiniPure Goat Anti-Rabbit IgG (Jackson ImmunoResearch Laboratories) secondary antibody at a dilution of 1:1000 in PBS containing 1% BSA.For nuclear counterstaining, DAPI (Servicebio) was used. After the staining process, slides were drained and mounted using AntiFade Mounting Medium (Servicebio) to preserve the fluorescence and prevent photobleaching. The stained tissue sections were visualized under a confocal microscope.

**Fractionation of terminal ileum to epithelial cell-enriched and immune cell-enriched fractions**

Mice aged 12-14 weeks were sacrificed for the study. After euthanizing the animals, the TI was carefully dissected, opened, and rinsed with cold deoxygenated PBS.

To isolate epithelial cells, the tissue was cut into segments and immersed in 1x Hank's Balanced Salt Solution (HBSS) supplemented with 20% fetal bovine serum (FBS), 1 mM dithiothreitol (DTT), and 2 mM ethylenediaminetetraacetic acid (EDTA). The segments were then incubated at

37 °C for 30 min with shaking to facilitate cell dissociation. The resulting solution was passed through a fine mesh filter to remove debris, and the suspended cells were pelleted by centrifugation at 212 x g. The cells were subsequently washed twice with PBS.

For the isolation of immune cells, the remaining tissue was immersed in PBS containing 0.492 mM $CaCl_2$ and 0.9 mM $MgCl_2$, supplemented with 20% FBS, collagenase (1 mg/ml, C0130, Sigma), and DNase (0.1 mg/ml, 4716728001, Sigma). Similar to the epithelial cell isolation, the segments were incubated at 37 °C for 30 min with shaking. The resulting solution was then passed through a fine mesh filter, and the suspended immune cells were pelleted by centrifugation at 212 x g and washed twice with PBS.

Both the isolated epithelial and immune cells were snap-frozen in liquid nitrogen and stored at -80 °C until further use. Tissue fractionation was carried out in an anaerobic environment to ensure the preservation of cellular integrity and oxygen-sensitive molecules. Notably, all solutions used for the isolation procedures were thoroughly degassed to minimize exposure to oxygen during cell isolation and maintain an anaerobic environment.

**Cultivation of cell lines**

Caco2 cells (ATCC CR2 2101) and Raw 264.7 cells (a kind gift of Prof. B. Levi, Technion) were cultured in DMEM (Sigma) supplemented with 10% heat-inactivated fetal calf serum (FCS) (Biological Industries), 1% L-glutamine (Biological Industries) and 1% Pen-Strep (Biological Industries), at 37°C in 5% $CO_2$. Periodic testing for mycoplasma contamination was conducted on both cell lines to ensure data reliability.

**Primary cultures of bone marrow-derived macrophages (BMDM)**

Femurs and tibias were harvested from C57BL/6 mice (WT and $Irp1^{-/-}$). Bone marrow cells were obtained by thorough flushing of the bones with DMEM. Following red blood cell lysis, cells were plated in DMEM supplemented with 20% FCS, 30% CCL1 cell-conditioned medium (CCL1 cells were a kind gift of Dr. J. Kaplan, University of Utah), 1% L-glutamine and 1% Pen-Strep (all from Biological Industries). The cells were cultured for 6 days in petri dishes.

On day 6, fully differentiated macrophages were scraped from the petri dishes on ice. The cells were then re-plated in PBS for 30 min. Subsequently, the PBS was replaced with DMEM containing 10% FCS, 1% Pen-Strep, and 1% glutamine for further experimentation.

**Inter-species co-culture assembly**

The day before co-culture assembly, Caco2 cells were grown in an oxygen-modified incubator (6% $O_2$). Confluent co-cultures of epithelial cells and macrophages were obtained by seeding 2 $x10^5$ cells per $cm^2$ surface area. For Caco2-Raw 264.7 co-cultures, Raw 264.7 cells were seeded and then Caco2 cells were added to them after 24 h. For Caco2-BMDM co-cultures, on day 6 after bone marrow extraction, the cells were plated as described above, and Caco2 cells were added to them after 24 h. Co-cultures were cultivated in an oxygen-modified incubator (6% $O_2$), at 37°C and 5% $CO_2$. Treatments of ferric ammonium citrate (Ammonium iron (III) citrate (FAC), F5879, Sigma), desferrioxamine (deferoxamine mesylate salt (DFO), D9533, Sigma), lipopolysaccharide (from *Escherichia coli* O111:B4, L4391, Sigma) and Tempol (4-Hydroxy-TEMPO (TEMPOL), 176141, Sigma) were carried out as detailed in the figure legends.

**Cell lysate preparation**
After the specified treatments, cells were lysed in degassed lysis buffer comprised of 10 mM HEPES (pH 7.2), 3 mM $MgCl_2$, 40 mM KCl, 5% glycerol, 0.2% Nonidet P-40, 5 mM DTT, 1 mM AEBSF (4-(2-Aminoethyl) benzenesulfonyl fluoride hydrochloride), 10 g/ml leupeptin and Complete$^{TM}$ EDTA-free protease inhibitor cocktail (11836170001, Roche). The lysis process was carried out for 10 min on ice. Nuclei and cellular debris were removed by centrifugation at 4$^{o}$C for 15 min at 14,500 *x g*. Cell lysates were prepared in an anaerobic chamber to maintain oxygen-freee environment. The protein concentrations in the cell lysate were determined using the Bradford assay.

**Western blotting and antibodies**
Equal amounts of protein (40 µg/lane) were separated by SDS-PAGE and transferred onto nitrocellulose or PVDF membranes. The transferred membranes were blocked with a blocking solution and probed with primary antibodies at 4 °C, as detailed in Table S1. Secondary horse-radish peroxidase-conjugated IgG antibodies used are listed in Table S3. Blots were developed using enhanced chemiluminescence (ECL kit, Pierce) to visualize the proteins bands on the membranes.

**RNA mobility shift assays**
Gel retardation assays were performed following the previously described method [4]. Cell lysates were prepared as described above-mentioned section. 5 µg total protein were added to a final volume of 12.5 µl buffer containing 25 mM Tris-HCl (pH 7.5) and 40 mM KCl, with or without 2% 2-mercaptoethanol (ME), which activates IRP1 *in vitro*. The protein samples were incubated for 5 min at RT with 12.5 µl reaction cocktail containing 20% glycerol, 0.2 U/µl Super RNAsine (Ambion), 0.6 µg/µl yeast tRNA (J61215, Thermo Scientific), 5 mM DTT and 20 nM $^{32}$P-labelled (NEG508H250UC, Perkin Elmer) IRE from the human ferritin H chain gene. The reaction was conducted in 25 mM Tris-HCl (pH 7.5), and 40 mM KCl. After the incubation, Samples (20 µl) of the reaction were loaded onto a 10% acrylamide/TBE gel. The gel was run at 200 V for 2.25 h. Following electrophoresis, the gel was then fixed, dried, and exposed for autoradiography.

**Aconitase assays**
Mitochondrial and cytosolic aconitase activities in cell lysates were assayed concurrently after electrophoretic separation, as previously described protocol [5]. Protein concentrations in the cell lysate were quantified using the Bradford assay 400 µg of protein were loaded into 8% polyacrylamide/TBE gel supplemented with 1 M Na citrate and run at 180 V for 5 h in running buffer [25 mM tris base, 192 mM glycine, and 3.6 mM citrate (pH 8)]. Gel was washed with distilled water and incubated at 37°C for 5 to 45 min in reaction solution [100 mM tris (pH 8.0), 1 mM nicotinamide adenine dinucleotide phosphate, 2.5 mM cis-aconitate, 5 mM $MgCl_2$, 1.2 mM MTT (3-(4,5-dimethylthiazol-2-yl)-2,5-diphenyltetrazolium bromide), 0.3 mM phenazine methosulfate, and isocitrate dehydrogenase (5 U/ml)]. All chemicals were purchased from Sigma. Gel is washed three times with distilled water for 5 min to remove background and gel is imaged.

**Quantitative RT-PCR**
Total RNA was extracted using the Trizol reagent (Invitrogen). For cell samples, cells were harvested directly with Trizol, while for tissue samples, whereas tissues were first ground in liquid $N_2$-cooled mortars and then incubated with Trizol, in accordance with the manufacturer's

instructions. Isolated RNA samples were subjected to DNase treatment using the DNase I recombinant, RNase-free kit (Quanta biosciences). Complementary DNA (cDNA) was synthesized using the cDNA Reverse Transcription Kit (Quanta biosciences). RT-qPCRs were performed using the SYBR master mix (Applied Biosystems). All primers used in the study are listed in Table S5. The relative expression of the target genes was calculated using the delta Ct method, normalizing to the expression of topoisomerase 1 for epithelial cells and ubiquitin for macrophage genes.

**Subcellular fractionation**

Cells were first suspended in lysis buffer comprised of 10 mM HEPES (pH 7.9), 10 mM KCl, 1.5 mM $MgCl_2$, 0.34 M sucrose, 10% glycerol, 1 mM DTT, 1 mM AEBSF and Complete TM EDTA-free protease inhibitor cocktail. Triton 0.1% was then added to the cell suspension and incubated for 5 min. The cell suspension was then centrifuged at 1500 × *g* for 4 min. The supernatant was carefully transferred to a new tube and was considered the crude cytoplasm-enriched fraction. The pellet obtained from previous step was then washed with the first lysis buffer then lysed in a second lysis buffer comprised of 3 mM EDTA, 0.2 mM EGTA, 1 mM DTT, 1 mM AEBSF and Complete TM EDTA-free protease inhibitor cocktail. The lysate was incubated for 10 min and then centrifuged at 1700 × *g* for 4 min. The supernatant was transferred to a new tube and was considered the nucleoplasmic fraction. The pellet obtained from previous step was then first washed in the second lysis buffer then suspended in the first lysis buffer supplemented with 1 mM $CaCl_2$ and 0.6 U of MNase (NEB-M0247S, New England Biologicals). The sample was incubated for 30 min at 37 °C). After centrifuging the sample at 20,000 × *g* for 10 min, the supernatant was transferred to a new tube and was considered the chromatin-bound fraction. The crude cytosol-enriched fraction was further centrifuged at 20,000 × *g* for 10 min, the supernatant was transferred to a new tube and was considered the cytosol-enriched fraction. All steps were performed in an anaerobic environment, and all lysis buffers were degassed.

**Labeling cells with $^{55}$Fe**

To detect iron inside the cells and in media, cells were given medium containing transferrin loaded with labeled iron. 200 mM of NTA (Nitrilotriacetic acid) dissolved in 0.9% NaCl and equilibrate to pH 3 with NaOH (10N). 100 mM cold $FeCl_3$ ($Fe^{56}$) was dissolved in 0.1M HCl. Hot- $Fe(NTA)_2$ was prepared in a volume ratio of 4:1:5 by mixing cold $FeCl_3$, hot $Fe^{55}Cl_3$ ($Fe^{55}$ NEZ-043-001MC, Perkin Elmer), and NTA, respectively. The mixture was incubated for 30 min at RT.
Subsequently, to allow for iron loading, apotransferrin (T1428, Sigma, in HBSS) was mixed with hot-$Fe^{55}(NTA)_2$ at a molar ratio of 1:4, comprising 125 µl of 1 mM apotransferrin, 12.5 µl of 0.5 M $NaHCO_3$, and 10 µl of 50 mM hot $Fe^{55}(NTA)_2$. Following a short incubation, which yielded a pink solution, 2 ml of medium was introduced. The resultant mixture was then filtered and added to the cells. After overnight incubation, the medium was removed, and cells were washed twice with PBS. To induce inflammation in the $^{55}$Fe labeled co-cultures, LPS (200 ng/ml) was added for the indicated time intervals. Harvested cells were snap-frozen and stored at -80°C until further use.

**Metabolic labeling and immunoprecipitation**

Cells were metabolically labeled for 1 h with $^{35}$S translabel (NEG772007MC, Perkin Elmer, mCi per 5 ml DMEM) without cysteine and methionine. The medium was supplemented with 10% dialyzed FCS, 10 mM HEPES, 1% pen/strep and 1% glutamine. After metabolic labeling, cells were lysed in lysis buffer comprised of 1% Triton X-100, 20 mM Tris (pH 8.0), 37 mM NaCl,

10% glycerol and EDTA-free protease inhibitor cocktail. Lysates were first pre-cleared with non-coated protein A-sepharose beads (L00210, GeneScript). The supernatants from the pre-cleared samples were then transferred to fresh tubes containing slurry of protein A-sepharose beads coated with anti-FtH antibodies (GE-Healthcare). The samples were rotated for 3 h at 4 $^{o}$C and then centrifuged at 800 × *g* to isolate the immunoprecipitated proteins. The immunoprecipitated proteins were denatured with reducing sample buffer x1 and boiled at 95 ℃ for 5 min. Subsequently, samples were then resolved by 14% SDS–PAGE. The SDS-PAGE gel was fixed, dried, and exposed for autoradiography to visualize the labeled proteins.

**S-nitroso-N-acetylpenicillamine (SNAP) treatment**

The day before SNAP treatment, Caco2 cells were cultured in MEM (Sigma) supplemented with 20% heat-inactivated FCS, 20 μM FAC, 1mM non-essential amino acids (Gibco), 10mM sodium pyruvate (Gibco), 2mM L-glutamine, and 1% Pen-strep. The cells were cultured in an oxygen-controlled incubator (6% $O_2$) at 37°C with 5% $CO_2$.

SNAP (N3398, Sigma) was added to the cells for 22 h, administered in two sequential doses. The first dose of SNAP was added to the cells, and they were incubated for 6 h. After the initial 6-hour incubation, 20% of the medium was collected from the culture and replaced with medium containing 5 mM SNAP. The cells were then further incubated for an additional 15 h.

SNAP Treatment in $^{55}$Fe-labeled Caco2 Cells:

Caco2 cells were treated with SNAP following the same protocol as described above. After the overnight SNAP incubation, cells were harvested at each designated time point. Subsequently, the cells were snap-frozen and stored at -80°C for further analysis.

**Griess assays**

Nitrite concentration in the medium was determined using the Griess assay. Briefly, the medium was collected and centrifuged at 1000 × *g*. Supernatant (50 μl) was then added to 50 μl of 1% sulfanilamide in 5% phosphoric acid. The mixture was incubated in the dark for 10 min. Next, 50 μl of 0.1 N-1-naphthylethylenediamine dihydrochloride (NED) was added. The reaction product was measured with the spectrocolorimeter (Multiskan EX, Thermo Electron Corporation), at 540 nm. $NaNO_2$ was used to create a standard curve of nitrite.

**Ferritin transfection**

HEK293T cells were grown in DMEM supplemented with 10% heat-inactivated FBS, 2 mM L-glutamine and 1% penicillin-streptomycin. The indicated ferritin constructs were transfected with polyethylenimine, according to standard protocol. Cells were harvested ~40 h post-transfection for subsequent analysis.

**Native-PAGE stains**

Samples were heat treated for 10 min in 70$^{o}$C and precipitated before separation on 6% Native-PAGE to visualize ferritin complexes.

**Perls'-DAB iron stain**

Gel is rinsed and then incubated for 1 h at RT with Perls' Prussian blue staining solutions (4% potassium ferrocyanide and 1.3% (v/v) HCl, mixed right before incubation). Gel is rinsed until neutral pH achieved and incubated in 100 mM Tris buffer (pH 7), for 5 min. The stain is then enhanced by incubation with 3,3′-Diaminutesobenzidine (DAB, 328005000, Thermo Scientific) solution (1%DMSO, 0.05% $H_2O_2$, 0.025% DAB in 100mM Tris buffer, pH 7) for 2 h for initial signal and overnight for saturation.

**Coomassie stain**

After electrophoresis, native gels were gently washed in the tap water and placed directly into staining solution InstantBlue® Coomassie Protein Stain (Ab119211, Abcam), according to the manufacturer's protocol.

**Human terminal ileum tissues**

Human biopsies were obtained from patients with clinically active Crohn's disease (CD) who were undergoing routine endoscopy or colonoscopy as part of their standard medical care. CD diagnosis was based on standard clinical, radiological, endoscopic, and histologic criteria. Biopsies were collected from both, inflamed and normal-appearing mucosa segments of each patient. The study protocol was approved by the institutional committee (IRB number 0052-17-RMB) and each patient provided informed consent before participating in the study.
Inflammation in the biopsied tissues was determined based on the endoscopic appearance of the mucosa. Inflammation was characterized (among others) by ulceration, erosion, lack of vascularity, edema, and erythema. Paraffin-embedded blocks of the human TI segments used for this study were provided by the Rambam Medical Center histology unit. Immunofluorescent staining of ferritin was performed following the same protocol as described for mouse samples.

**Statistical analysis**

Statistical analysis was performed using Graph Pad Prism 8 (GraphPad Software). Values in graphs are expressed as mean ± SD (unless otherwise specified). Data were examined for normality of distribution and variance homogeneity. Two groups of mice were compared with Student's 2-tailed t test for independent data. $p < 0.05$ (*$p < 0.05$, **$p < 0.01$,***$p < 0.001$, ****$p < 0.0001$) was considered significant.

**Table S1: Experimental details of Western blot analysis of tissue lysates – 1st antibody**

| Antigen | Lower gel (%) | Blocking solution | 1st Antibody | Dilution | Solution | Temp. | Time |
|---|---|---|---|---|---|---|---|
| ACTIN | 12% | 5% BSA | Santacruz, sc-1616 | 1:2500 | 5% BSA | RT | 1 h |
| TFR1 | 12% | 5% BSA | Abcam, cat# ab84036 | 1:2000 | 5% BSA | 4 ºC | ON |
| IRP2 | 12% | 5% milk | a kind gift of Dr. Tracey Rouault | 1:2000 | 5% milk | 4 ºC | ON |

BSA: bovine serum albumin; ON: overnight; RT: room temperature

**Table S2: Experimental details of Western blot analysis of cell culture lysates – 1st antibody**

| Antigen | Lower gel (%) | Blocking solution | 1st Anti-body | Dilution | Solution | Temp. (ºC) | Time |
|---|---|---|---|---|---|---|---|
| ACTIN | 12% | 5% BSA | sc-1616 | 1:10,000 | 5% BSA | RT | 1h |
| TFR1 | 6% | 5% BSA | Invitrogen 136800 | 1:500 | TBST | 4ºC | ON |
| IRP2 | 12% | 5% milk | a kind gift of Dr. Tracey Rouault | 1:200 | 5% milk | 4ºC | ON |
| FTH1 | 12% | 2% milk | Cell signaling D1D14 | 1:1000 | 2% milk | 4ºC | ON |
| FTL | 12% | 1% milk | Ab 69090 | 1:1000 | 1% milk | 4ºC | ON |
| HIF2α | 10% | 5% milk | NB100-122b | 1:500 | 5% milk | 4ºC | ON |
| ERK 1/2 | 12% | 5% BSA | sc-514302 | 1:1500 | 1% BSA | 4ºC | ON |
| p-ERK 1/2 | 12% | 5% BSA | sc-7383 | 1:1500 | 1% BSA | 4ºC | ON |
| p-NFκB 65 | 12% | 5% milk | sc-136548 | 1:500 | 1% BSA | 4ºC | ON |
| p38 | 12% | 5% milk | sc-81621 | 1:500 | 1% BSA | 4ºC | ON |
| p-p38 | 12% | 5% milk | sc-7973 | 1:500 | 1% BSA | 4ºC | ON |

BSA: bovine serum albumin; ON: overnight; RT: room temperature

**Table S3: Experimental details of Western blot analysis of cell culture lysates - 2nd antibody**

| 2nd Antibody | Dilution | Solution | Temp. | Time |
|---|---|---|---|---|
| Horseradish-peroxidase conjugated anti goat (Sigma A5420) | 1:10,000 | TBST | RT | 1h |
| Horseradish-peroxidase conjugated anti mouse (Danyel biotech) | 1:10,000 | TBST | RT | 1h |
| Horseradish-peroxidase conjugated goat anti rabbit (Abcam) (Abcam ab97200) | 1:25,000 | TBST/<br>5% milk (HIF2α) | RT | 1h |

TBST: 1X Tris-Buffered Saline, 0.1% Tween® 20 Detergent; RT: overnight

**Table S4: Sequence of primers used for murine genes in quantitative PCR analysis**

| Primer name | Primer sequence |
|---|---|
| *Ubc* Forward | CCA GTG TTA CCA CCA AGA AG |
| *Ubc* Reverse | ACC CAA GAA CAA GCA CAA GG |
| *Tnfα* Forward | CAA ATT CGA GTG ACA AGC CTG |
| *Tnfα* Reverse | GAG ATC CAT GCC GTT GGC |
| *Icam1* Forward | CAA TTT CTC ATG CCG CAC AG |
| *Icam1* Reverse | AGC TGG AAG ATC GAA AGT CCG |
| *Il1b* Forward | GCC CAT CCT CTG TGA CTC AT |
| *Il1b* Reverse | AGG CCA CAG GTA TTT TGT CG |
| *Il6* Forward | AGC CAG AGT CCT TCA GAG AGA T |
| *Il6* Reverse | TGT TAG GAG AGC ATT GAG AAT TGG |
| *Tfr1* Forward | GTG GAG TAT CAC TTC CTG TCG |
| *Tfr1* Reverse | CCC CAG AAG ATA TGT CGG AAA GG |
| *Dmt1* Forward | CAT CCT GGT CCT GAT CGT CTGC |
| *Dmt1* Reverse | CCA CAT AGA GTG CCA CAT GCC CTAG |
| *Fpn1*+IRE Forward | AGG CTT TGG CTT CCA ACT TCA GC |
| *Fpn1*+IRE Reverse | AAG CAC AAC AGC CTT ATG CCG AAA G |
| *Fth1* Forward | AAG TGC GCC AGA ACT ACC AC |
| *Fth1* Reverse | TTC AGA GCC ACA TCA TCT CG |
| *Ftl1* Forward | AAT CAG GCC CTC TTG GAT CT |
| *Fth1* Reverse | GGT TGC CCA TCT TCT TGA TG |
| *Ncoa4* Forward | AAG AAA GTG GGA AGC CTC AG |
| *Ncoa4* Reverse | AGA TCA CAA ACT GCT GGG AG |
| *Fxn* Forward | TAC ATG TCA CAG TCA ACG CC |
| *Fxn* Reverse | CTC GTC TAG AGA GCT TGG GT |
| *Hsc20* Forward | AAT GAA AGA CTC GCA GAC GC |
| *Hsc20* Reverse | AAG TCA CCT TGT TCA AAA GCG |
| *Iscu* Forward | GAT TGT GGA CGC CAG ATT CA |
| *Iscu* Reverse | GGC TTC CTC CAC CGT TTT C |
| *Lyrm4* Forward | ATA ATC CGC AGA CAG GTC CA |
| *Lyrm4* Reverse | TTC CTT GTG CTG GCT TCC TA |

#### Table S5: Sequence of primers used for human genes in quantitative PCR analysis

| Primer name | Primer sequence |
|---|---|
| *TOP1* Forward | GGT GAG AAG GAC TGG CAG AAA T |
| *TOP1* Reverse | CTT GTC GAT GAA GTA CAG GGC TA |
| *IL8* Forward | ATGAGTTCCAAGCTGGCCGTGGCT |
| *IL8* Reverse | TCTCAGCCCTCTTCAAAAACTTCT |
| *CXCL10* Forward | CTG CCA TTC TGA TTT GCT GC |
| *CXCL10* Reverse | CGT ACA GTT CTA GAG AGA GG |
| *ICAM1* Forward | TTG GGC ATA GAG ACC CCG TT |
| *ICAM1* Reverse | GCA CAT TGC TCA GTT CAT ACA CC |
| h*TFR1* Forward | TGG AGA CTT TGG ATC GGT TGG TG |
| h*TFR1* Reverse | CAG TGG CTG GCA GAA ACC TTG |
| *DMT1* Forward | CAC CAT GAC AGG AAC CTA TTC TGG C |
| *DMT1* Reverse | AAC CAC TCG GGC AAA GCG TGA C |
| *FPN1*-IRE Forward | AAA CGA AGT CAA CCA AGG CTA CAG TC |
| *FPN1*-IRE Reverse | ACA GGA GTG CAA GGA ACT GGA GAT AG |
| *FTH1* Forward | AAA CGC GAA TTC ACC ATG ACG ACC GCG |
| *FTH1* Reverse | ACC GCG GCC GCT TAG CTT TCA TTA TCA |
| *FTL* Forward | GAA CCA GGC CCT TTT GGA TC |
| *FTL* Reverse | CCA GGA AGT CAC AGA GAT GGG |
| *NCOA4* Forward | CAA ATT CCT GAG CAC TTG ATG G |
| *NCOA4* Reverse | AAA GGG ACA GCT ACA ATA CCG |
| *FXN* Forward | ACT AGC AGA GGA AAC GCT GG |
| *FXN* Reverse | GGA ATA GGC CAA GGA AGA CA |
| *NFS1* Forward | ACT CCC GGA CAC ATG CTT ATG |
| *NFS1* Reverse | GCT GAC GAG CAC GTT CCAT |
| *NARF* Forward | GGC GCT GAC TGC GTG TTA A |
| *NARF* Reverse | AGG TCA CCT TGC TCC ATT ATT TG |
| *LYRM4* Forward | AAA GCC AAG AGA GAC CTT GGA |
| *LYRM4*Reverse | CTA GGT CCT GGG CAT GTC TC |
